# Multicomponent Thiolactone-Based Ionizable Lipid Screening Platform for an Efficient and Tunable mRNA Delivery to the Lungs

**DOI:** 10.1101/2024.10.09.617396

**Authors:** Álvaro Peña, Juan Heredero, Beatriz Blandín, Elena Mata, Diego De Miguel, Alfonso Toro, Teresa Alejo, Diego Casabona, Alexandre López, Ana Gallego-Lleyda, Esther Pérez-Herrán, Juan Martínez-Oliván, Javier Giménez-Warren

**Affiliations:** Certest Pharma, Certest Biotec S. L., 50840, San Mateo de Gállego (Zaragoza), Spain

**Keywords:** lipid nanoparticle, ionizable lipid, drug delivery, lipids, extrahepatic, lung targeting, mRNA therapeutics

## Abstract

Ionizable lipids are an essential component of lipid nanoparticles (LNPs) for an efficient mRNA delivery. However, optimizing their chemical structures for high protein expression, efficient endosomal escape, and selective organ targeting remains challenging due to complex structure-activity relationships and multistep synthesis. In this study, we introduce a rapid, high-throughput platform for screening ionizable lipids using a two-step, scalable synthesis involving a one-pot 3-component click-like reaction. This method, herein known as the STAAR approach, standing for Sequential Thiolactone Amine Acrylate Reaction, allowed for the combinatorial synthesis and in vivo screening of 91 novel lipids, followed by a structure-activity study. This led to the development of CP-LC-0729, an ionizable lipid that significantly surpasses the benchmark in protein expression while showing no in vivo toxicity. Additionally, the STAAR lipid platform was further validated by incorporating a one-step strategy to yield a permanently cationic lipid which was tested following a fifth-lipid formulation strategy. The in vivo results showed a highly selective lung delivery with a 32-fold increase in protein expression, outperforming current endogenous targeting strategies. All these findings underscore the potential of lipid CP-LC-0729 and the STAAR lipid platform in advancing the efficiency and specificity of mRNA delivery systems, while also advancing the development of new ionizable lipids.

## Introduction

The interest in mRNA technology as a promising platform for the treatment of diverse pathologies, including cancer, has increased exponentially since the COVID-19 pandemic, when two mRNA vaccine candidates (i.e. mRNA-1273 and BNT162b) were authorized to tackle SARS-CoV-2. Indeed, mRNA-based technology has shown great potential in applications such as gene editing therapies, vaccines against infectious diseases and protein supplementation therapies.^1–3^ In order to achieve protein expression in living organisms, mRNA molecules need a delivery platform to avoid nuclease degradation, promote cellular endocytosis and improve endosomal escape. Lipid nanoparticles (LNPs) demonstrate both advanced clinical status and effectiveness in overcoming delivery barriers for mRNA therapeutics, highlighting their importance as nonviral platforms.^4,5^

LNPs commonly consist of mRNA, helper lipids (usually phospholipids), cholesterol, PEG-conjugated lipids and ionizable lipids. The role of ionizable lipids has been demonstrated as crucial in stabilizing and delivering mRNA.^6,7^ A fundamental chemical feature of ionizable lipids lies in the tertiary amino group which becomes charged in acidic pH but remains neutral at physiological conditions^8^. During the formulation process at acidic pH, their positive charge aids in condensing the mRNA and, upon internalization, as the endosomes undergo acidic maturation, they assist in membrane disruption, facilitating endosomal escape. At physiological pH, ionizable lipids exhibit a neutral charge, resulting in reduced toxicity levels and enhanced biocompatibility.^9,10^ Two substantial research challenges currently being addressed involve understanding and improving the efficiency of LNPs’ endosomal escape (currently estimated to be around 2%) and achieving targeted in vivo delivery to specific cells or tissues for precision therapeutics.^11,12^

Despite the extensive research and advances into ionizable lipid development, ionizable lipids typically consist of distinct common structural components: an ionizable headgroup and at least two hydrophobic alkyl tails tethered together through a linker, often containing degradable bonds such as esters. Ester bonds have been chosen for their ability to break down in vivo thanks to the activity of esterases, which improves their biocompatibility.^13,14^ Moreover, ionizable lipids commonly include at least two hydrophobic tails, which can foster a cone-like shape, improving endosomal escape and mRNA delivery by promoting the formation of inverted hexagonal phases.^12,15^ Adjusting the bonds and chemical structures in ionizable lipids to achieve effective and safe in vivo delivery has sparked interest recently. However, three significant challenges persist regarding ionizable lipid design and synthesis: Firstly, despite some general characteristics previously outlined, precise structural design rules are still lacking regarding the relationship between their structural features and in vivo delivery.^9^ Moreover, major bottlenecks in mRNA delivery, such as endosomal escape, could be addressed by gaining a better understanding of structure-activity relationships.^12^ Indeed, the most recurrent approach to novel ionizable lipid development is to conduct massive screenings of synthetic candidates based on combinatorial chemistry to identify those that are particularly effective in drug delivery.^16–18^

Secondly, addressing the necessity for screening lipid candidates requires avoiding multi-step and complicated synthetic protocols, highlighting the importance of rapid and cost-effective synthetic strategies.^19,20^ Numerous strategies have been explored so far, for example, aza-Michael addition with acrylates, ester formation, and epoxide opening.^16^ However, obtaining the necessary building blocks typically involves several time-consuming syntheses, harsh conditions and purification steps, which can hinder structural optimization and once again the understanding of structure-activity relationships.^13,21^

Thirdly, there is a growing interest in developing LNPs that can target certain tissues or organs through passive targeting.^11^ In this sense, one of the most successful strategies up to date is the inclusion of a fifth lipid in the composition of the LNPs. Siegwart and coworkers developed formulations by incorporating either cationic or anionic lipids, thereby enabling targeted delivery to the lungs or spleen, respectively.^22^

Here, we developed a rapid, high-throughput and cost-effective platform for screening ionizable lipids. We utilized a versatile multidimensional synthesis approach involving mild conditions and two tandem one-pot reactions: thiolactone ring opening and Michael addition. We then investigated the structure-activity relationships of hydrophobic tails and functional groups leading to the best ionizable lipid candidate. Finally, the combination of this candidate, CP-LC-0729, with a permanently cationic lipid, which can be accessed via a single synthetic step, demonstrated a very high selectivity to lungs with a remarkable 32-fold increase in protein expression compared to the benchmark. Altogether, these results highlight the potential of the STAAR platform in generating new promising lipid candidates for both hepatic and extrahepatic therapeutic applications.

## Results and Discussion

### Sequential Thiolactone Amine Acrylate Reaction (STAAR Lipids)

Similarly to other ionizable lipid structures, STAAR lipids consist of a polar head, a linker, and hydrophobic tails. Given that one of the aims of this study was precisely to examine how the various fragments of these lipids can influence their *in vivo* performance, a combinatorial chemistry approach was followed to screen a selection of lipids that included different moieties in their structure. For simplicity, each lipid was named as AiBjCk to account for the reagents used for their synthesis. In this nomenclature, Ai represents an amine (polar head); Bj indicates the linker, which consists of a thiolactone incorporating a hydrophobic tail, and Ck represents an acrylate with another hydrophobic tail.

The synthetic process of STAAR lipids is illustrated in Figure 1a. Briefly, this two-step synthesis starts with the coupling of homocysteine thiolactone with a carboxylic acid to yield the corresponding linker intermediate (Bj). The second step involves a tricomponent one-pot sequential reaction which starts with a ring-opening addition of the intermediate (Bj) with an amine (Ai) followed by a Michael addition with an acrylate (Ck) to yield the lipid AiBjCk^23^. Remarkably, this multicomponent reaction can be carried out at room temperature, without need for catalysts or inert atmosphere, and within a 2-hour reaction time. Moreover, the high purity (>80%) obtained makes it an optimal method for the rapid multidimensional screening of ionizable lipids, eliminating the need for further purification steps. This method simplifies the synthesis of new ionizable lipids in 1.5 mL tubes, reducing the reliance on expert organic synthesis knowledge, which allowed the rapid screening of nearly 100 novel lipids.

**Figure 1.**
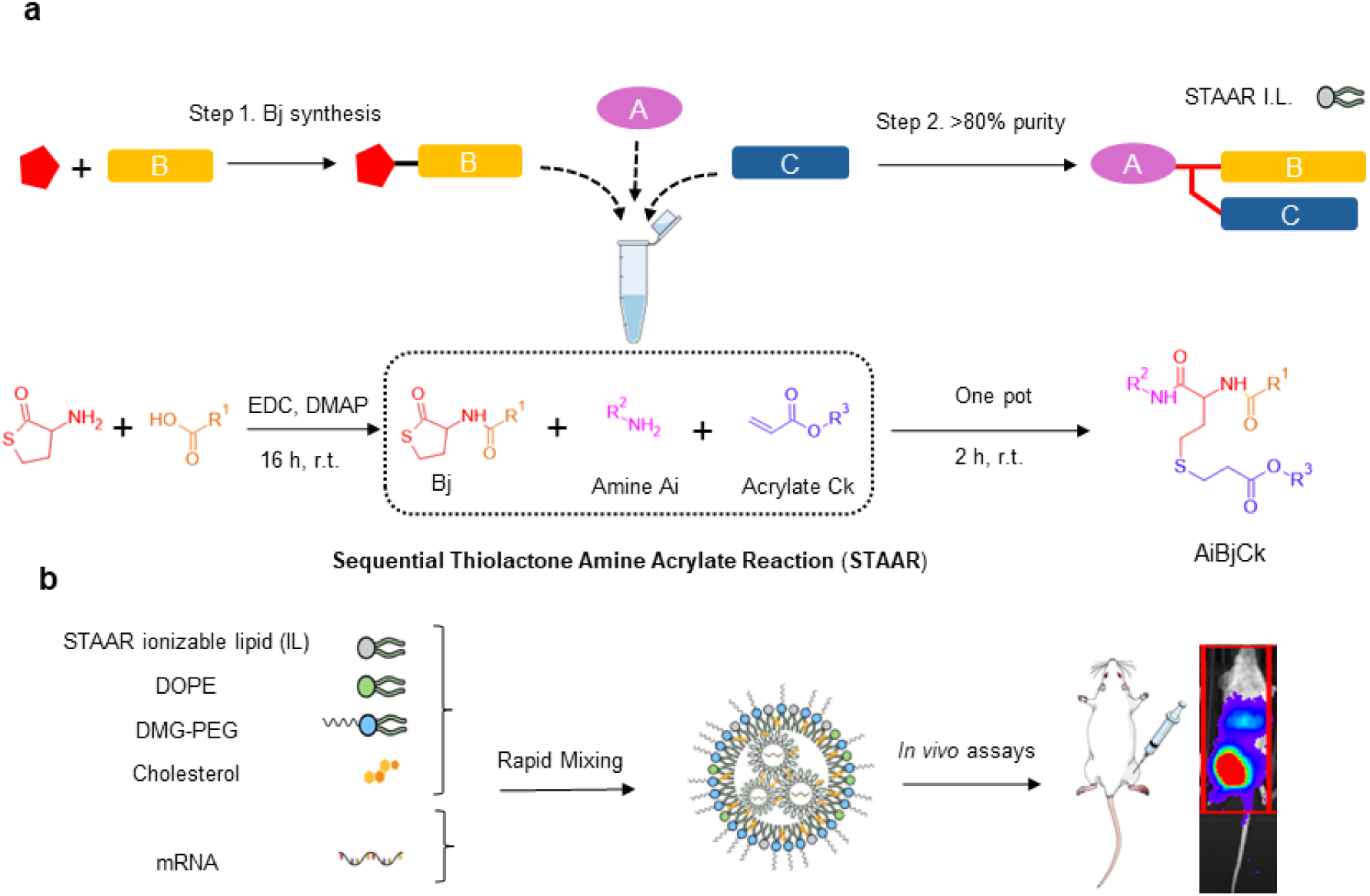
Design and synthesis scheme of STAAR ionizable lipids. a Representative scheme describing STAAR lipids synthetic steps. First, the synthesis of Bj thiolactone derivative and then the one-pot tricomponent reaction. The amine opens the thiolactone ring of Bj (aminolysis) and subsequently the thiol group reacts with the acrylate to afford the final lipid with high purity in short times and at room temperature.^23^ b Scheme of the process followed for the formulation of lipids into LNPs for mRNA delivery *in vivo*.

The general structure of the resulting STAAR lipids includes two amide bonds (one originating from the reaction of the amine Ai and the thiolactone Bj and the other one connecting this thiolactone to its hydrophobic tail), a thioether bond (from Bj), and an ester group (from Ck). Consequently, any changes to be made in the hydrophobic tails can be achieved by modifying the structure of the thiolactone derivatives (Bj) and acrylates (Ck).

### Two-phase screening strategy for ionizable lipids

To assess how different moieties affect the final structure and activity of the lipid, a two-phase screening strategy was devised in which one component of the lipid (Ai, Bj, or Ck) was initially kept constant while the other two were varied. This method enabled us to systematically evaluate the impact of each component on the lipid’s performance. By fixing one fragment and varying the other two, the intention was to efficiently identify the most promising candidates, thereby streamlining the screening process and optimizing our approach. Thus, the screening process was divided into two optimization phases, with each phase focusing on the optimization of a different lipid moiety (Figure 2a, 2b). This methodological approach not only reduced the number of required experimental animals but also enhanced the efficiency of the screening process.

**Figure 2.**
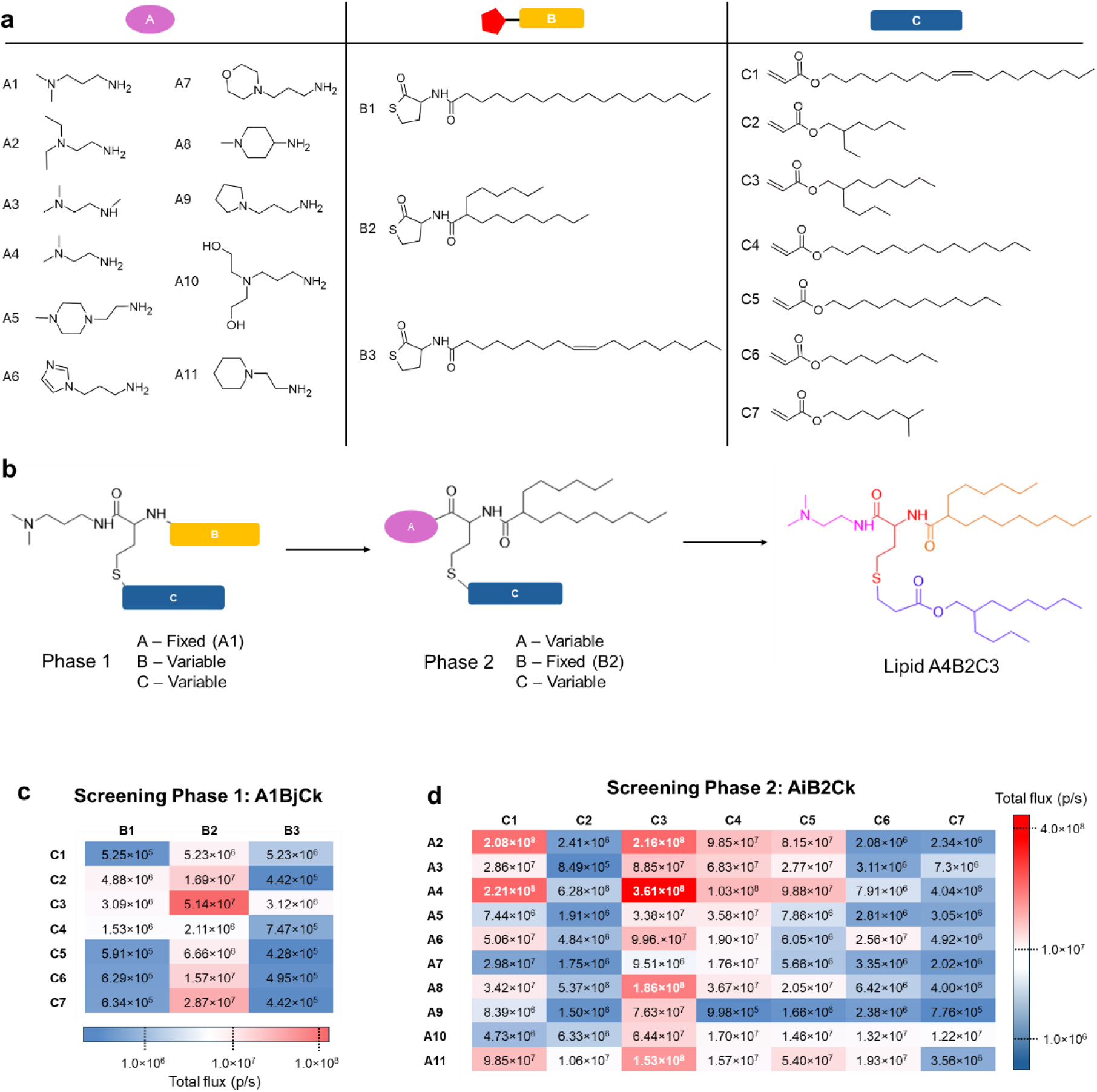
Screening optimization of STAAR lipids. a Amines (Ai), Bj derivatives and acrylates (Ck) used for lipid screening. b Scheme summarizing the strategy followed for the two screening phases. c Total flux results of screening phase 1, where B was optimized by combining different acrylates and Bj derivatives while keeping polar head (A1) fixed. Mice were intramuscularly injected (i.m.) with mRNA-Luc-loaded LNPs at an mRNA dose of 0.05 mg/kg. Luminescence imaging acquisition was performed at 4 h post-treatment and total flux was quantified. *In vivo* mLuc expression displayed in a heat map (n= 3 biologically independent samples). Data are presented as mean values. d Total flux results table of Screening phase 2 where A and C were optimized by combining different amines and acrylates while keeping B2 fixed. Mice were i.m injected with mRNA-Luc-loaded LNPs at an mRNA dose of 0.05 mg/kg. Total flux luminescence imaging acquisition was performed at 4 h post-treatment and total flux was quantified. *In vivo* mLuc expression displayed in a heat map (n= 3 biologically independent samples). Data are presented as mean values.

For this optimization phase, the effectiveness of each IL candidate was evaluated by formulating each lipid individually into LNPs using a pipette rapid mixing procedure.^24^ Thus, each LNP was composed of an ionizable lipid (AiBjCk), 1,2-dioleoyl-sn-glycero-3-phosphoethanolamine (DOPE), cholesterol, and 1,2-dimyristoyl-rac-glycero-3-methoxypolyethylene glycol-2000 (DMG-PEG-2000) (50: 10: 38.5: 1.5 molar ratios, respectively) and firefly luciferase mRNA (with a 6:1 molar N/P ratio Figure 1b).

Also, due to the weak correlation between *in vitro* and *in vivo* LNP performance,^25^ a direct screening *in vivo* using mice as animal model was followed, for which a firefly luciferase mRNA dose of 0.05 mg/kg was administered intramuscularly (i.m.) and the luminiscence results were measured at 4h post-inoculation.

### Screening Phase 1: Optimization of linker’s hydrophobic chain (Bj)

The first phase of the structure optimization, aimed to select the optimal Bj component by synthesizing 21 lipids in which the polar head was fixed (A1 was selected) while combining three types of thiolactones with different acrylates (Figure 2c). This amine was chosen as its structure closely resembles that of the MC3 polar head, featuring a spacer of three methylene units between a tertiary amine and a carbonyl group.^26^ The hydrophobic chains linked to the thiolactone derivatives (Bj) were selected to have distinctly different structures: a linear chain (B1) with 18 carbon atoms, an unsaturated chain (B2), and a branched chain (B3).

The analysis of the protein expression (Figure 2c) achieved for the LNPs for each individual lipid revealed that A1B2Ck lipids (with k ranging from 1 to 7) resulted in increased protein expression for *in vivo* mRNA delivery, regardless of the acrylate used (Ck). This observation aligns with existing literature, which suggests that branched structures often confer enhanced performance in lipid-based delivery systems.^27^ Moreover, unsaturated chains yielded higher expression than saturated ones. Notably, hydrophobic moieties C3 and C7 were particularly effective.

### Screening Phase 2: Optimization of polar head (Ai) and second hydrophobic chain (Ck)

Based on the findings from the screening phase 1, B2 was selected as the foundation for the next screening step. This second phase aimed to optimize the other two moieties of the molecule: the polar head (Ai) and the second hydrophobic chain (derived from Ck). Once again, the use of the STAAR synthesis approach for this optimization process allowed for a swift exploration of these structural moieties thanks to the high purities achieved in the sequential reactions used in their synthesis. More specifically, 70 AiB2Ck novel lipids were synthesized, with i and k ranging from 2 to 11 and 1 to 7, respectively (Figure 2d). Moreover, to validate the strategy employed, an assay was carried out to corroborate that there were no significant differences between the use of crude reaction and purified ionizable lipid in terms of protein expression (Figure S3).

Regarding the Ai moiety, diverse polar heads, including cyclic, linear, aromatic, and analogues with a variable number of ionizable N groups, underwent testing. Upon *in vivo* evaluation, cyclic and aromatic amines (A5, A6, A7, A8, A9) failed to demonstrate significant expression, the only exception being that of A11. In contrast, among the linear amines, A2 and A4 displayed the best results, with lipid A4B2C3 notably yielding the highest expression levels. Remarkably, the comparison of amines A1 and A4, which differ by just one single methylene unit, resulted in an almost 5-fold increase in expression for lipid A4B2C3 over A1B2C3, emphasizing the role of the number of methylenes between the tertiary amine and the primary amine of Ai in the lipid performance. These observations showcase the sensitivity of lipid performance to minor structural changes in the spacer length of the polar head. Remarkably, the comparison of amines A1 and A4, which differ by just one single methylene unit, resulted in an almost 5-fold increase in expression for lipid A4B2C3 over A1B2C3, emphasizing the role of the number of methylenes between the tertiary amine and the primary amine of Ai in the lipid performance.^28^ These observations showcase the sensitivity of lipid performance to minor structural changes in the spacer length of the polar head performance.^28^ These observations showcase the sensitivity of lipid performance to minor structural changes in the spacer length of the polar head.

With regards to the hydrophobic tails (stemming from Ck), we confirmed the previously observed trend for Bj, with the butyloctanoate derived branched acrylate (C3) yielding the lipids with highest expression levels. Additionally, linear 18-carbon monounsaturated (C1) also showed promising results *in vivo*. We also noticed that when shorter acrylate chains were used, the *in vivo* expression decreased up to 60 times, irrespective of it being branched or linear, this including isooctyl (C7), ethylhexyl (C2), and linear octyl (C6). Moreover, intermediate length chains (C5 and C4, 12 and 14 carbons in length, respectively), led to higher expression compared to shorter chains, yet the expression remained three times lower than C3 and C1. This suggests that longer chains, featuring unsaturations or branches in the hydrophobic moiety Ck, might be necessary to achieve the desired electronic and steric balance in the STAAR lipids.

Altogether, this screening phase highlighted the profound impact of the C moiety, where branched and unsaturated Ck chains, especially C3, demonstrated substantial improvements in protein expression.

In relation to physicochemical parameters of LNPs, observed sizes ranged from 150 to 230 nm, with the majority falling beneath 190 nm. Polydispersity indexes mostly lay below 0.25, while ζ potentials ranged between 5 mV and -25 mV, with all lipids but four exhibiting a negative ζ potential. mRNA encapsulation rates varied from 20% to 100% (Table S2). Notably, lipids with higher activity achieved encapsulation rates between 60% and 80% (with a direct pipetting method), hence suggesting this might represent the optimal encapsulation range for STAAR lipids LNPs in the screening phase. Moreover, we noted that even though employing an amine featuring two ionizable groups (A5) as lipid head results in high encapsulation rate, this does not reflect in its performance, as the *in vivo* activity was found to be 2 to 30 times lower compared to lipids containing amine A4, which contains only one ionizable group.

### Fine-tuning of the ionizable lipid structure

Following the discovery of lipid A4B2C3 as the top-performing candidate in the initial two-phase screening approach, the research focused on exploring the impact of further modifications to the lipid structure, specifically targeting the hydrophobic branched chains.

After identifying A4B2C3 as the most effective ionizable lipid in the two-phase screening, further structural studies were conducted to assess the impact of hydrophobic chain lengths and functional groups and their roles in influencing lipid performance. The purpose of this stage was to deepen the understanding of how variations in the chemical structure can influence protein expression, thereby refining the structure-activity relationship for improved delivery efficiency.

### Assessment of the hydrophobic tails of lipid A4B2C3

Following the same synthetic approach as in the screening phase, we synthesized and purified eight additional ionizable lipids, hence introducing three distinct branched chain lengths via the Bj and Ck moieties (Figure 3a). Having optimized the lipid structure in the initial phases, for these refining phases we relied on microfluidic techniques to confirm the previous conclusions while keeping the formulation process precisely controlled. Nanoparticles were obtained with sizes ranging from 80 to 130 nm, with the majority being under 100 nm, and they exhibited high encapsulation efficiencies (90-99%, Table S4). Thus, a purified batch of the optimized lipid A4B2C3 along with these new 8 lipids were formulated into LNPs along with two separate controls. Even though both controls used MC3 as ionizable lipid in the same molar and N/P ratios as the screened lipids, one of them included DOPE as helper lipid (same as the formulated STAAR lipids), namely MC3 DOPE, whereas the other one replicated the MC3 LNP benchmark by using DSPC. Remarkably, when comparing the *in vivo* results to those of the MC3 benchmark LNP^29^ (Table S3), eight out of the nine tested lipids showed a total fux that was at least double that of the MC3 benchmark.

**Figure 3.**
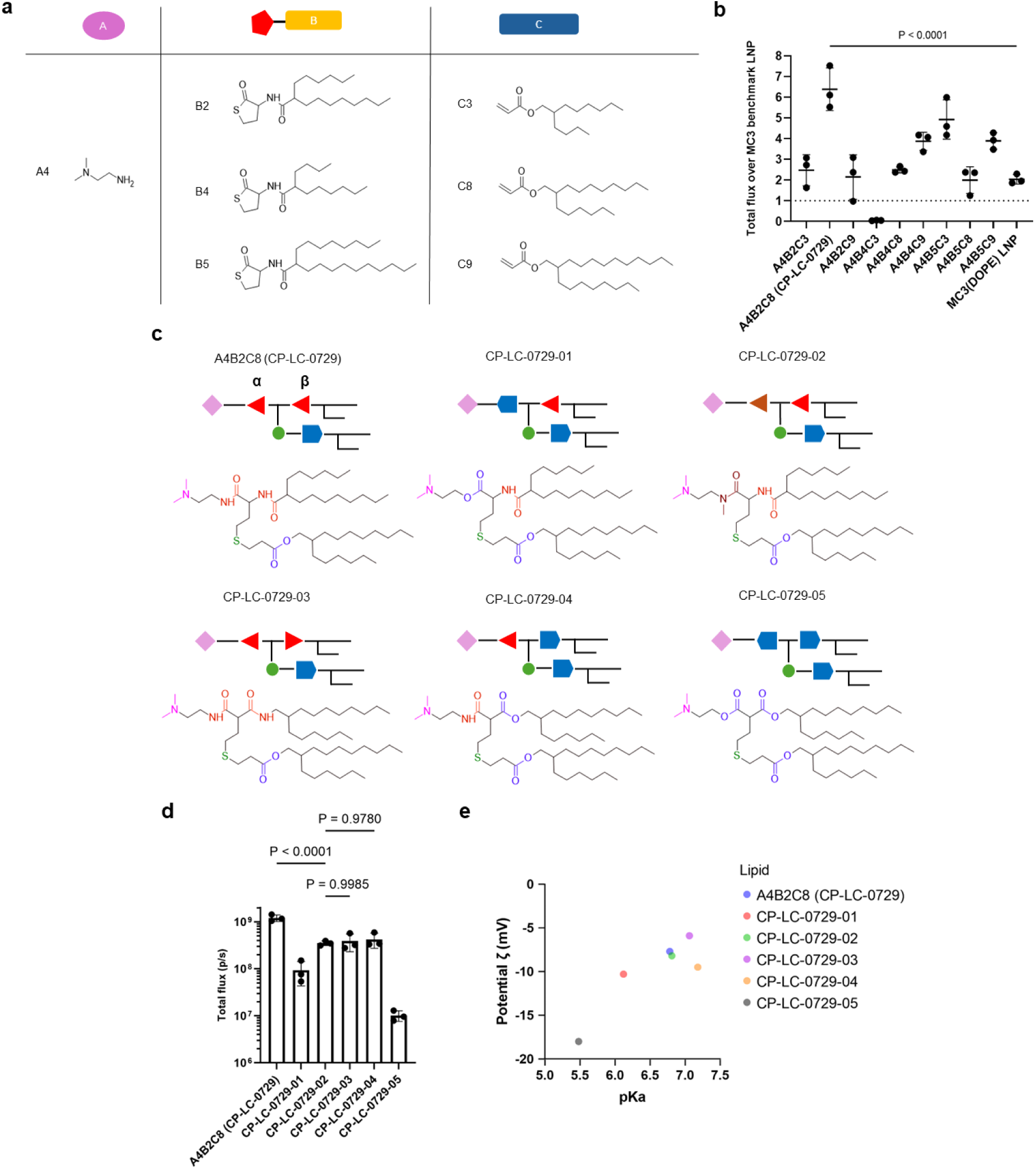
Assessment of the hydrophobic tails of A4B2C3 and structure-activity relationships in STAAR lipids. a Building blocks for hydrophobic tail optimization, combining A4, branched Bj derivatives and branched acrylates. b *In vivo* luminiscence total flux results of Figure 3a structures normalized respect to MC3 benchmark LNP, with the dashed line in the y-axis representing the results of control MC3 benchmark LNP. MC3(DOPE) LNP was also used as control employing DOPE as helper lipid. Mice were i.m injected with mRNA-Luc-loaded LNPs at an mRNA dose of 0.05 mg/kg. Luminescence imaging acquisition was performed at 4 h post-treatment and total flux was quantified. *In vivo* mLuc expression (n= 3 biologically independent samples). Data are presented as mean values ± standard deviation (SD).c Schematic lipids set structure displaying different functional groups varying in alpha (α) and beta (β) positions. d Results of lipids in Figure 3c, *in vivo* mLuc expression (n= 3 biologically independent samples). Data are presented as mean values ± SD. Mice were i.m injected with mRNA-Luc-loaded LNPs at an mRNA dose of 0.05 mg/kg. Luminescence imaging acquisition was performed at 4 h post-treatment and total flux was quantified. e ζ potentials and pKa of the formulated LNPs (n= 3 biologically independent samples). Data are presented as mean values. In b and d one-way ANOVA with Tukey’s correction was used.

Among these, the result for A4B2C8 (known as CP-LC-0729) was particularly striking, as it demonstrated an expression value 6.4 times higher than the benchmark control (Figure 3b), which firmly establishes this lipid as the best candidate for further studies and optimization. The exception amongst these 8 lipids was observed with lipid A4B4C3, characterized by the shortest hydrophobic chain combination, and for which the expression dropped to 0.034 times relative to the MC3 benchmark. The results suggest that a tail with a minimum carbon number may be necessary for achieving high protein expression, aligning with the trend observed during the two-phase screening, where shorter chains such as C2, C6, and C7 consistently resulted in lower protein levels.^19^

### Influence of functional groups in the structure-activity relationship of STAAR lipids

After exploring the impact of altering the hydrophobic chains and identifying lipid CP-LC-0729, which contains two branched chains (B2, C8), as the best candidate for high-performing *in vivo* activity, our focus then shifted to study the functional groups within the structure. As previously reported in literature, certain bonds, such as esters, play a pivotal role in endosomal escape and degradation pathways.^7,14,28,30^ Taking advantage of the chemical versatility of STAAR lipids, which allows for modular modification of functional groups, we investigated how the nature of specific bonds can influence the performance of the STAAR lipids.

We focused specifically on the role of the amide bonds, which are common to all previously synthesized STAAR lipids and are located at positions referred to as α and β,^31,32^ by synthesizing purified structural variations of CP-LC-0729 as depicted in Figure 3c. These products were subsequently formulated with the same N/P and lipid ratios used in the initial screenings, employing microfluidic mixing.

As a first step, we inverted the C-N carbonyl order (N-CO to CO-N) of the β amide bond (CP-LC-0729-03) resulting in a threefold reduction of *in vivo* protein expression compared to CP-LC-0729 (Figure 3c and 3d). This change may be attributed to the shift in the relative position of the β amide hydrogen bond, which can potentially affect the intermolecular force landscape and packing of the molecule.^33^

Then, to further explore the role of the α amide bond, lipid CP-LC-0729-02 was synthesized, where the α amide is methylated, resulting in a 3.4-fold decrease of *in vivo* expression. In the next step, the amide bonds were partially or completely substituted with ester bonds in CP-LC-0729-01, CP-LC-0729-04 and CP-LC-0729-05. In this case, we noticed a significant decrease in effectiveness—by two orders of magnitude—when replacing both amides (α and β positions) with ester groups in CP-LC-0729-05 (Figure 3d). This change resulted in the lowest performance among the proposed variations, and contrasts with the fact that other commercial ionizable lipids such as ALC-0315, SM-102, or MC3 only feature ester groups. As a middle-ground case, when combining an amide and an ester group, the *in vivo* performance was dependent on relative position of the groups, yielding intermediate outcomes. Interestingly, when lipids contain only one amide, their expression is four and a half times higher when located in the α position (CP-LC-0729-04) compared to the β position (CP-LC-0729-01). Thus, these results indicated that preserving the α amide bond (CP-LC-0729-04) led to better expression results than replacing it with an ester group (CP-LC-0729-05). Remarkably, the protein expression value of the methylated α amide lipid CP-LC-0729-02 was on par with CP-LC-0729-03 and CP-LC-0729-04. Interestingly, when lipids contain only one amide, their expression is four and a half times higher when located in the α position (CP-LC-0729-04) compared to the β position (CP-LC-0729-01). Additionally, in the case of the methylated α amide (CP-LC-0729-02), which no longer acts as a hydrogen bond donor, the expression decreased threefold compared to the non-methylated counterpart, suggesting that hydrogen bonds might play a key role. We hypothesize that the observed difference may arise from the α amide proximity to the polar head, potentially enhancing the pKa and its availability to interact with mRNA via hydrogen bonding or intermolecular interactions between ionizable lipids. The optimal combination, as evidenced by our findings, involves amides located at both the α and β sites, reflecting in the chemical structure of lipid CP-LC-0729.

Regarding the LNP physicochemical characterization, we noticed that altering the α amide bond led to lower LNP pKa. Specifically, pKa values were the lowest for CP-LC-0729-05 (pKa=5.48) and CP-LC-0729-01 (pKa =6.12). A similar trend was observed when measuring ζ potential, as CP-LC-0729-05 (ζ=-18.0 mV) and CP-LC-0729-01 (ζ=-10.3 mV) displayed the lowest values of the series (Figure 3e).

Additionally, to further evaluate the behavior of lipid CP-LC-0729 in different formulations, an *in vivo* study using LNPs with different molar ratios of the four lipid components was conducted. The results, detailed in Table S7, indicate that CP-LC-0729 performs well even when the molar ratios diverge from the ones used in more conventional formulations, highlighting its versatility.

### Additional biological assays of the top-performing ionizable lipid CP-LC-0729 LNP

### *In vivo* safety profile of lipid CP-LC-0729

After selecting CP-LC-0729 as the best performing ionizable lipid candidate through the *in vivo* screening and subsequent optimization phases, different biological assays were carried out to evaluate the safety profile. Thus, *in vivo* toxicity assays on CP-LC-0729 LNPs were conducted to assess any potential organ damage by measuring liver enzymes (ALT, AST, ALP) and kidney function (UREA) at 24 and 48h after systemic injection at a high dose (2.5 mg/kg). When compared to the PBS buffer used as a control, CP-LC-0729 LNPs showed no significant impact on the levels of these enzymes (Figure 4a). The change in mice body weight was monitored for 15 days post-inoculation, revealing no apparent differences compared to the negative control (PBS buffer) (Figure 4b). While a more comprehensive assessment, including testing across different animal models, is necessary to establish a complete safety profile of this lipid, our preliminary results indicate excellent tolerability following systemic administration, with no signs of toxicity observed under the conditions studied.

**Figure 4.**
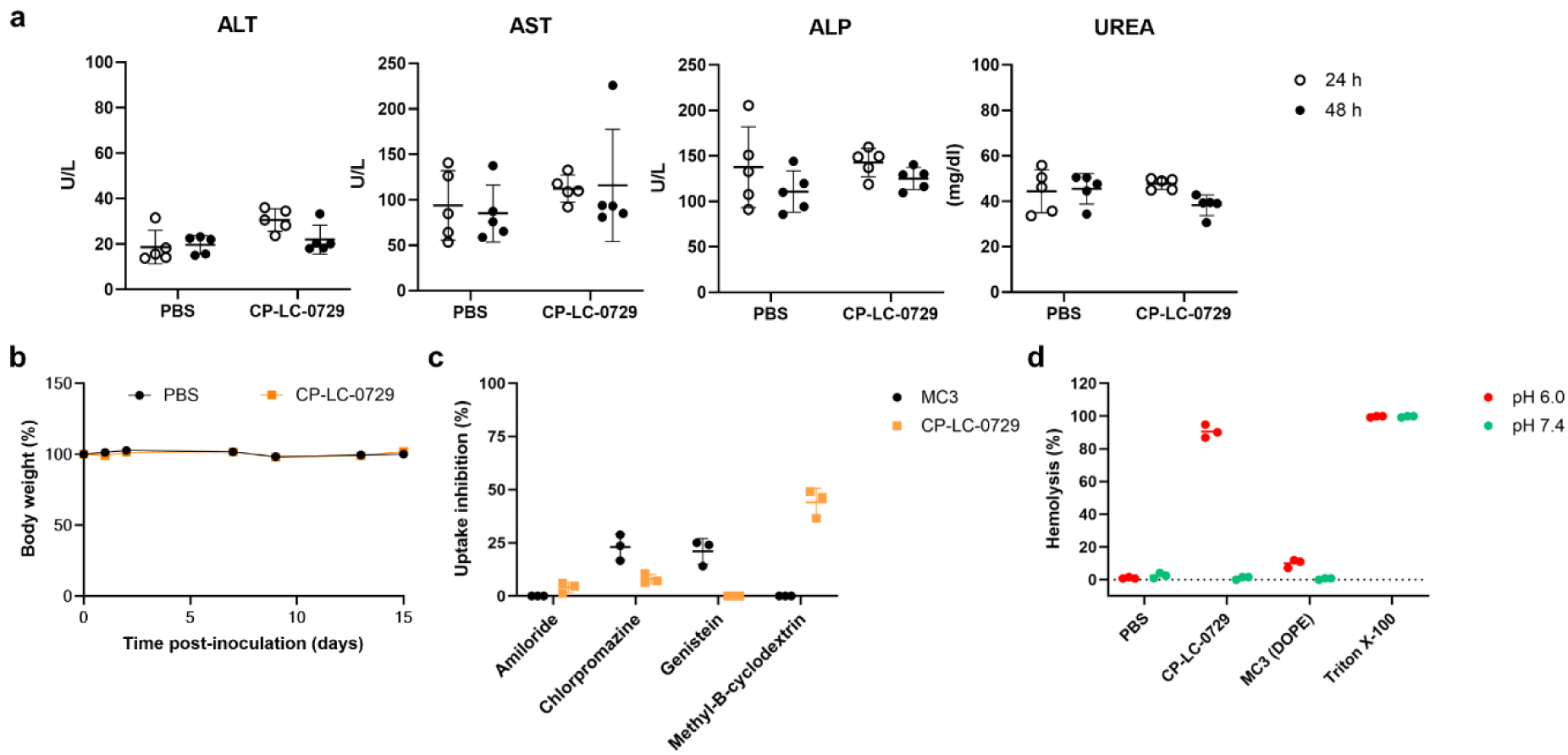
*In vivo* safety and cell internalization profile of CP-LC-0729 LNP. a Serum alanine aminotransferase levels (ALT), serum aspartate aminotransferase levels (AST), serum alkaline phosphatase levels (ALP), urea nitrogen concentration (UREA). Mice were intravenously injected (i.v) with mRNA-Luc-loaded LNPs at an mRNA dose of 2.5 mg/kg (n= 5 biologically independent samples). Serum was collected for ALT, AST, ALP and UREA analysis at 24 h and 48h post-treatment. Data are presented as mean values ± SD. Two-way ANOVA with Tukey’s correction was used. b Body weight change for 15 days (measured made at days 0, 1, 2, 7, 9, 13 and 15). Mice were i.v injected with mRNA-Luc-loaded LNPs at an mRNA dose of 2.5 mg/kg (n= 5 biologically independent samples). Data are presented as mean values ± SD. c Uptake inhibition of CP-LC-0729 and MC3(DOPE) LNP by different endocytosis inhibitors (n = 3 biologically independent samples). The inhibitors used were Amiloride (inhibitor of macropinocytosis), Chlorpromazine (inhibitor of clathrin-mediated endocytosis), Genistein (inhibitor of caveolae mediated endocytosis), Methyl-β-cyclodextrin (inhibitor of lipid raft mediated endocytosis). Data are presented as mean values ± SD. d Hemolysis assay of CP-LC-0729 LNP and MC3(DOPE) LNP at pH 7.4 or 6.0. RBCs were incubated with CP-LC-0729 LNP and MC3(DOPE) LNP at an mRNA concentration of 2,5 μg/mL at 37 °C for 1 h. Positive and negative controls were carried out with 0.1% Triton-X and 1× PBS, respectively. Data are presented as mean ± SD (n = 3 biologically independent samples).

### Evaluation of cell internalization of CP-LC-0729 LNPs

To gain deeper insights into the mechanisms of cell internalization, the endocytosis pathways of CP-LC-0729 LNP were examined *in vitro* in hepatic HepG2 cells. Four endocytic pathways were assessed by incubating the LNPs with specific inhibitors for each route before *in vitro* transfection^19^. The results show that the uptake of CP-LC-0729 LNPs strongly relies on lipid rafts, as indicated by inhibition with methyl-β-cyclodextrin. Other internalization pathways, such as macropinocytosis and clathrin-mediated endocytosis, showed a smaller contribution. Additionally, genistein did not inhibit uptake, indicating that caveolae-mediated endocytosis is not significant (Figure 4c). We also conducted hemolysis assays to assess endosomal escape of CP-LC-0729 LNPs through their membrane-disrupting activity under both acidic (pH 6) and physiological (pH 7.4) conditions.^19,27^ These tests showed that CP-LC-0729 LNPs increased hemolysis in the acidic medium up to 90%, ten times higher than the MC3 (DOPE) LNP used as control. In contrast, at physiological pH, the hemolytic activity remained comparable to the control level (PBS) indicating no hemotoxicity. (Figure 4d) This suggests a potential enhanced endosomal escape mechanism of CP-LC-0729 LNPs at acidic pH, while maintaining hemocompatibility at physiological conditions, a result that is in accordance with the fact that lipid CP-LC-0729 shows a higher in vivo performance than MC3 while displaying no signs of toxicity.^34^

### Extrahepatic organ targeting of CP-LC-0729 using permanently cationic lipids

Recognizing the critical importance of targeting specific organs for effective therapeutic outcomes, we focused on achieving extrahepatic mRNA delivery to broaden the scope and versatility of ionizable lipid CP-LC-0729 and the STAAR screening platform. As reported by Siegwart and coworkers,^22,35,36^ the addition of a fifth permanently charged lipid results in selective targeting of the lungs or spleen, attributed to changes in the protein corona and LNP surface charge^11^. Herein we explored a streamlined approach to modify the STAAR ionizable lipid CP-LC-0729 into its permanently cationic form (+) CP-LC-0729, using a single synthetic step, and later carried out a comparative study with what is reported in the literature following the fifth-lipid strategy. Hence, a novel cationic lipid was synthesized, denoted as (+) CP-LC-0729, as it is derived from the methylation of the tertiary amine of the top-performing ionizable lipid CP-LC-0729 (Figure 5a). Subsequently, the performance of ionizable lipids MC3 and CP-LC-0729 was compared, in LNPs incorporating cationic lipids (+) CP-LC-0729 or the standard commercial alternative DOTAP as fifth lipids, using representative formulations with firefly luciferase mRNA (Figure 5b).^37^ As shown in Table S5, the cationic lipid molar percentages were compared using LNPs that incorporated different proportions of this fifth lipid (30%, 40% and 50% (mol/mol)). The experiments were conducted using microfluidics mixing for LNPs, followed by intravenous administration. LNPs were termed as [cationic lipid (percentage of cationic lipid in molar rate) ionizable lipid] LNP, for example, LNP using DOTAP as cationic lipid at 50% combined with MC3 as ionizable lipid was designated [DOTAP (50%) MC3] LNP.

**Figure 5.**
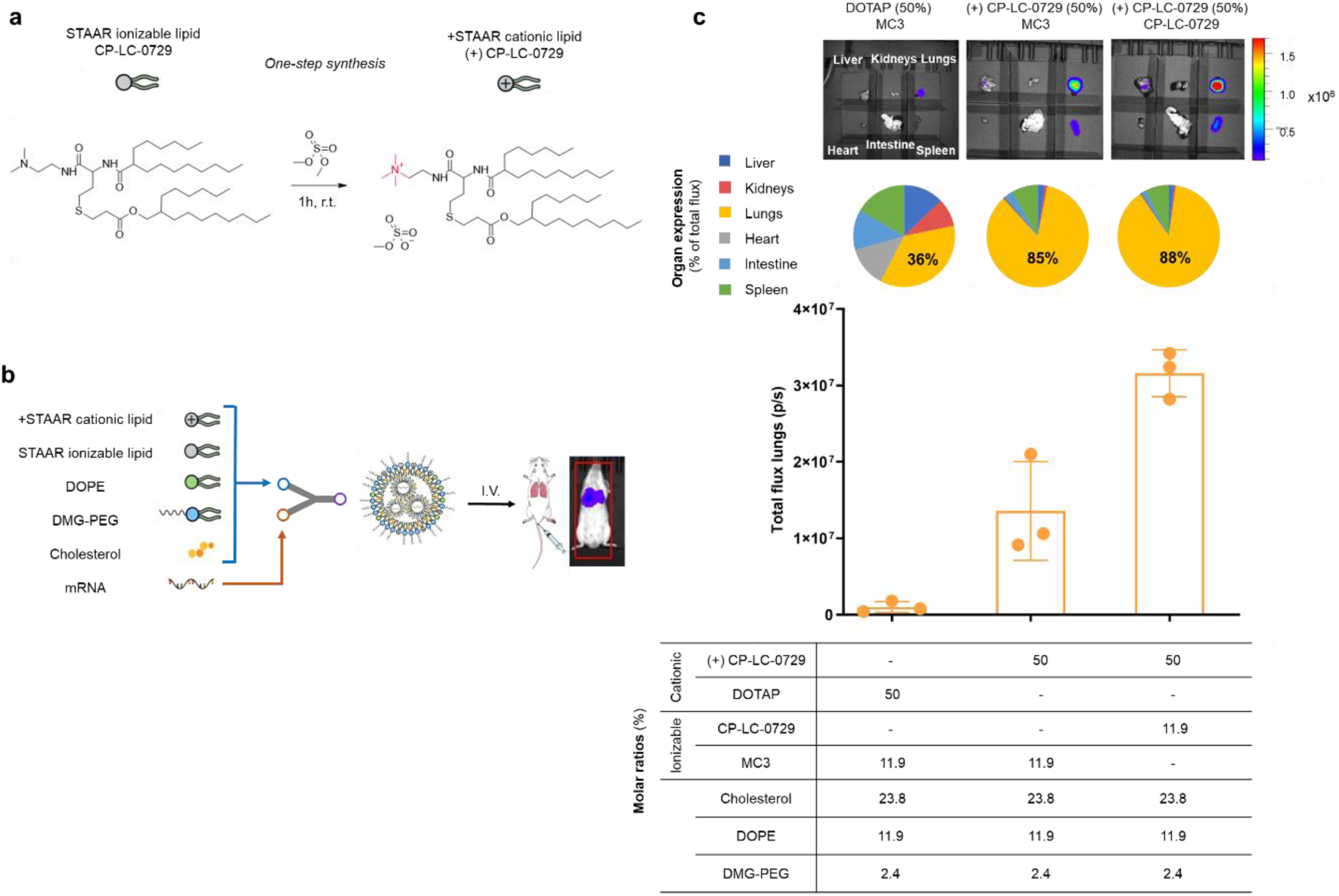
Lung targeting strategy using a STAAR permanently cationic lipid. a Synthetic scheme used to prepare a permanent cationic lipid, namely (+) CP-LC-0729, from an ionizable lipid CP-LC-0729. b Schematic representation of the formulated LNPs incorporating a STAAR permanently cationic lipid as fifth component for mRNA delivery *in vivo* through i.v. administration. c Top to bottom: *Ex vivo* luminescence of major organs from mice, pie charts showing a profile of the tissue-specificity of mLuc expression in the lungs, heart, liver, intestine, spleen and kidneys, bar representation of total luminescence found in lungs and table with the formulation details of LNPs (n= 3 biologically independent samples). Mice were i.v. injected with mRNA-Luc-loaded LNPs at an mRNA dose of 0.2 mg/kg. Images were acquired at 4h post-treatment. For the bar graph, data are presented as mean values ± SD. One-way ANOVA with Tukey’s correction was used.

Regarding the physiochemical properties, LNPs with (+) CP-LC-0729 showed similar values to CP-LC-0729 LNP, including particle size (60-85 nm) and 99 % mRNA encapsulation efficiency. However, we noticed positive surface ζ potentials in (+) CP-LC-0729 LNPs, which varied depending on the cationic lipid ratio used (Table S6).

After inoculation, *ex vivo* assays revealed distinct variations in protein expression in the lungs, depending on the cationic lipid ratio. The [(+) CP-LC-0729 (50%) CP-LC-0729] lipid nanoparticles (LNPs) demonstrated remarkable selectivity for the lungs, achieving 88% selectivity. Even the [(+) CP-LC-0729 (40%) CP-LC-0729] and [(+) CP-LC-0729 (30%) CP-LC-0729] LNP exhibited strong selectivity, 88% and 82%, respectively (Fig S4 and S5), far exceeding the 38% selectivity observed for the benchmark LNP, [DOTAP (50%) MC3].

Moreover, when [(+) CP-LC-0729] was formulated with MC3, the resulting expression levels were twofold lower than when it was combined with CP-LC-0729 (Figure 5c). Even more remarkably is the fact [(+) CP-LC-0729 (50%) CP-LC-0729] LNP exhibited 32-fold higher luciferase expression compared to the benchmark [DOTAP (50%) MC3] LNP (Figure 5c). These findings demonstrate that both the selectivity and protein expression of CP-LC-0729 combinations significantly outperform the benchmark [DOTAP (50%) MC3] formulation. Furthermore, the optimal results observed when combining (+) CP-LC-0729 with CP-LC-0729 may suggest that structural similarity between the ionizable and cationic forms may improve lipid packing and enhance overall LNP performance. However, more studies are needed to confirm this hypothesis. This result shows the potential of CP-LC-0729, while also expanding its application to therapies requiring targeted extrahepatic delivery.

Additionally, [(+) CP-LC-0729 (50%) CP-LC-0729] LNP was formulated with a small percentage of the fluorescent probe DiR (dialkylcarbocyanine, 1,1’-dioctadecyl-3,3,3’,3’-tetramethylindotricarbocyanine iodide) to track fluorescence and determine LNP biodistribution. Interestingly, while the majority of LNPs were directed to the liver, luciferase expression was observed predominantly in the lungs (Figure S6).

## Conclusion

The STAAR platform enables the synthesis of ionizable lipids in a high-throughput, one-pot setup, offering cost-efficiency, modular structure design and mild reaction conditions with shortened times which can easily be scaled up. Through the combinatorial synthesis and *in vivo* screening of 91 molecules, we identified several effective STAAR lipid candidates during two optimization phases. We then explored key structural characteristics governing their activity through structure-activity relationship studies, including hydrophobic tails and functional groups, identifying lipid CP-LC-0729 as the top-performing candidate and laying the groundwork for discovering new effective STAAR lipids in the future. Compared to the benchmark MC3, ionizable lipid CP-LC-0729 exhibited 6.4 times higher protein expression and showed no signs of *in vivo* toxicity, positioning it as a promising, readily scalable candidate for clinical advancement. We further validated the efficacy of the STAAR platform by introducing a one-step strategy for developing cationic lipids. The use of lipid CP-LC-0729, in combination with its permanently cationic counterpart, produced lipid nanoparticles (LNPs) that achieved a remarkable 32-fold increase in lung protein expression compared to the benchmark SORT LNP (MC3/DOTAP), with 88% organ selectivity. This result once again confirms the potential of CP-LC-0729, while also expanding its application to therapies requiring targeted delivery beyond the liver. Lastly, acknowledging the vast array of possibilities unlocked by the STAAR platform, ongoing studies aim to expand the scope and structures of this platform and explore the delivery mechanisms.

## Experimental section

### Materials

All chemical reagents were purchased from Sigma Aldrich (Burligton, Massachusetts, USA), Tokyo Chemical Industry (TCI, Tokyo, Japan), Fluorochem Ltd (Hadfield, UK) and Ambeed (Arligton Heights, Illinois, USA). 1,2-dioleoyl-sn-glycero-3-phosphoethanolamine (DOPE), 1,2-distearoyl-sn-glycero-3-phosphocholine (DSPC) were purchased from Avanti Polar Lipids (Alabaster, Alabama, USA). 1,2-dimyristoyl-rac-glycero-3-methoxypolyethylene glycol-2000 (DMG-PEG 2000) was purchased from Cayman Chemicals (Ann Arbor, Michigan, USA). Cholesterol was purchased from Sigma Aldrich (Burligton, Massachusetts, USA). DLin-MC3-DMA (MC3) was purchased from Broadpharm (San Diego, California, USA). DOTAP was purchased from BOC Science (New York, USA).

### mRNA Synthesis

Firefly luciferase mRNA was synthesized following a previous reported method^38^. mRNA complete sequence is available in Figure S1.

### General method for the synthesis of STAAR ionizable lipids

STAAR lipids were synthesized using a tandem one-pot reaction. First, thiolactone derivatives (0,15 mmol, 1 equiv.) and acrylate (0,15 mmol, 1 equiv.) were dissolved in 300 ul tetrahydrofuran (THF) at RT, followed by the addition of amine (0,15 mmol, 1 equiv.). Two hours later, THF was removed under vacuum and the resulting crudes were used for initial screening. All crudes were characterized by mass spectrometry and their purity was assessed by HPLC coupled with a light scattering detector (ELSD) (Table S1).

### Purification and characterization of ionizable lipid A4B2C8 (CP-LC-0729)

To purify the top-performing lipid, namely CP-LC-0729, its crude product was separated using a CombiFlash NextGen 300+ with gradient elution from 100% of dichloromethane to 50% of 80/20/1 DCM/MeOH/ NH_4_OH (aq). CP-LC-0729 was characterized by mass spectrometry (MS) and nuclear magnetic resonance spectroscopy (NMR). MS-QDa: theoretical [M+H]^+^ = 740.63, experimental [M+H]^+^=740.85; ^1^H NMR (400 MHz, CDCl_3_) δ: 6.76 (m, 1H); 6.37 (d, *J*= 7.9 Hz, 1H); 4.60 (q, J= 7.1 Hz, 1H); 3.98 (d, *J*= 5.8 Hz, 2H); 3.36 (m, 2H); 2.78 (t, *J*= 6.8 Hz, 2H); 2.59 (m, 4H); 2.48 (t, *J*= 5.9 Hz, 2H); 2.28 (s, 6H); 2.06 (m, 2H); 1.96 (m, 1H); 1.58 (m, 3H); 1.41 (m, 2H); 1.25 (m, 44H); 0.87 (m, 12H).

### STAAR cationic lipid (+) CP-LC-0729 synthesis

Lipid CP-LC-0729 (149.5 mg, 0.20 mmol) was dissolved in toluene (0.7 ml) under Argon atmosphere. Then, dimethylsulfate (22.9 ul, 0.24 mmol) was added and one hour later the solvent was evaporated under reduced pressure. Afterwards, the crude reaction was purified using a CombiFlash NextGen 300+ with gradient elution from 100% of DCM to 40% of MeOH to afford the pure product (70%).

MS-QDa: theoretical [M+H]^+^ = 754.66, experimental [M+H]^+^ = 754.84; ^1^H NMR (400 MHz, DMSO-d_6_) δ: 8.38 (m, 1 H); 8.17 (m, 1H); 4.29 (m, 1H); 3.92 (d, *J*= 5.6 Hz, 2H); 3.49 (m, 2H); 3.33 (m, 2H); 3.09 (s, 9H); 2.68 (m, 2H); 2.60 (m, 2H); 2.41 (m, 1H); 2.19 (m, 1H); 1.80 (m, 2H); 1.57 (m, 1H); 1.48 (m, 2H); 1.36-1.04 (m, 46H); 0.89 (m, 12H).

### Nanoparticle Formulation

#### General Procedure

For all LNP formulations, mRNA was dissolved in a pH 4, 10 mM sodium citrate solution and combined with the lipid mixtures, which comprised a combination of STAAR ionizable lipids, DOPE, cholesterol, and DMG-PEG2000, at an N/P ratio of 6. For the specific experiments including a fifth cationic lipid, these lipidic and N/P ratios were modified as detailed in Table S5.

In the initial screening process (optimization phases 1 and 2), a direct pipetting mixing method was used whereas a microfluidics equipment was employed for the subsequent phases and for the extrahepatic targeting formulations with the permanently cationic lipids. Hence, briefly, the methods followed in each of these stages is as follows:

Direct Pipetting Mixing (Lipid Screening Phases 1 and 2): The ethanolic lipid phase and the aqueous mRNA solution were mixed at an N/P ratio of 6 (mol/mol) by pipetting the aqueous solution into the organic phase with vigorous pipetting for about 10 seconds. The resulting LNPs were diluted 1:1 with a pH 8 buffered solution.^24^

Microfluidic Mixing (Fine-tuning of lipid structure and cationic lipid formulations): Both the ethanolic lipid phase and the aqueous mRNA solution were mixed using a NanoAssemblr^®^ Ignite™ (Precision NanoSystems) device with a flow rate ratio (FRR) of 3:1 and a total flow rate (TFR) of 12 ml/min. The resulting LNPs were dialyzed overnight against a pH 8 Tris buffer solution containing 15% sucrose. The final LNP solution was adjusted to a concentration of 100 µg/ml of mRNA, filtered through a 0.22 µm filter, and stored at 4°C.

### Characterization

^1^H-RMN were measured using a Bruker 400 MHz NMR spectrometer. HPLC-ELSD-MS was performed on a Waters Alliance equipped with Waters ELSD 2424 and Waters Acquity QDa detectors. The hydrodynamic size, zeta potential and polydispersity index (PDI) physicochemical parameters of LNPs were measured using a Malvern Zetasizer^®^ Advance Lab Blue Label Equipment. The mRNA encapsulation efficiency and the pKa of LNP were obtained using Quant-IT^®^ Ribogreen following the manufacturer’s instructions and a 6-(p-toluidinyl)naphthalene-2-sulfonic acid (TNS) assay, respectively.^39^ Characterization data is available in supporting information.

### Animals and cell lines

All animal experiments were conducted in agreement with European and national directives for protection of experimental animals. Experimental procedures were approved by the Ethics Committee for Animal Experiments of University of Zaragoza (PI07/23).

8-10 weeks old female BALB/cAnNRj mice were purchased from Janvier Labs. All mice were housed and maintained in specific pathogen–free conditions in the facilities of Centro de Investigaciones Biomédicas de Aragón (Zaragoza, Spain; reference ES 50 297 0012 01). Animals underwent a one-week acclimation period upon arrival at the research facilities. Housing conditions were controlled, maintaining a room temperature of 20-24 °C with 50%-70% humidity, a light intensity of 60 lux and a light-dark cycle of 12 hours. Food and water were provided *ad libitum*.

HepG2 cells were obtained from the American Type Culture Collection (ATCC). Cells were cultured on RPMI 1640 (Gibco, 31870074), supplemented with 10% Fetal Bovine Serum (Sigma, F7524), 1% Penicillin-Streptomycin Solution (Gibco, 15140122) and 2 mM Glutamax (Fisher, 35050038).

### Cellular internalization inhibition

HepG2 cells were seeded in Delta-treated 96-well plates at a density of 10.000 cells/well in complete DMEM and incubated overnight at 37 °C and 5 % CO_2_. Then, cells were pre-treated for 1h with endocytosis inhibitors: amiloride (1 mM), chlorpromazine (15 µM), genistein (150 µM) and methyl-B-cyclodextrin (1.25 mM). After the inhibition treatment, cells were transfected with LNPs encapsulating luciferase mRNA (200 ng mRNA/well) and formulated with the different ionizable lipids. After 4h incubation the cell medium was replaced with fresh complete DMEM and incubated overnight. Cells were lysed with PBS-Triton 0.1 %, cell lysate was transferred to an opaque 96-well white plate and d-Luciferin (GoldBio LUCK-100 resuspended in 100 mM Tris-HCl pH 7.8, 5 mM MgCl2, 250 μM CoA, 150 μM ATP buffer) was added to each well reaching a final concentration of 150 μg/mL. Luminescence was measured after 5 minutes of incubation at room temperature in a FLUOstar Omega plate reader (BMG Labtech). The efficiency of endocytosis was subsequently evaluated comparing luminescence signal of the untreated wells, and the wells treated with each inhibitor for each LNP.

### Hemolysis assay

Erythrocytes were isolated from fresh heparinized human blood sample. A volume of 2 mL of blood was washed three times with neutral PBS (pH 7.4) by centrifugation at 800g for 5 minutes. Afterwards, the sample was resuspended in a total volume of 1 mL either in neutral or acid PBS (pH 6). Isolated erythrocytes were diluted one hundred times in PBS with the corresponding pH. Erythrocytes were incubated with LNPs at a final mRNA concentration of 2.5 ug/ml at 37°C for 1 hour. Finally, the erythrocytes were centrifuged at 800g for 5 minutes and 60 μl of the supernatant was transferred to a 96-well plate. 0.1% Triton X-100 was included as a positive control for membrane disruption. Absorbance was measured at 540 nm in a FLUOstar Omega plate reader (BMG Labtech).^40^

### *In vivo* safety Evaluation

Animals were inoculated intravenously with 2.5 mg/kg of luciferase-encoding mRNA formulated in the indicated LNPs as described above. Throughout the experiment, mice were periodically weighed and monitored for observable clinical signs of toxicity.

Blood samples were collected from the submandibular vein on days 1 and 2 post-inoculation and serum samples were obtained for biochemical analysis. Samples were centrifuged at 8000 rcf at 4°C for 10 minutes and supernatants were collected. Biochemical analyses were performed using Cobas c-311 (Roche Diagnostics) according to manufacturer instructions.

### *In vivo* imaging

For intramuscular inoculation, animals were injected with 0.05 mg/kg of the indicated Luciferase-encoding mRNA-LNPs into the right thigh muscle. LNPs were diluted in Tris buffer containing 15 % sucrose in a final volume of 30 µL, and administered using a 30G insulin syringe.

For intravenous inoculation, animals were injected into the tail vein with 0.2 mg/kg of the indicated Luciferase-encoding mRNA-LNPs. LNPs were diluted in Tris buffer containing 15 % sucrose in a final volume of 250 µL and administered using a 27G needle.

At 4 hours post-inoculation, mice were anesthetized by inhalation of Isoflurane (IsoVet) 4% using an animal inhalation chamber. The maintenance of the anesthesia was sustained at 1.5% isoflurane. D-luciferin (12507, Quimigen) was intraperitoneally injected at 150 mg/kg diluted in PBS. Luminescence images were acquired 20 minutes after luciferin inoculation.

For *ex vivo* luminescence studies, mice were euthanized 20 minutes after luciferin injection and liver, spleen, lungs, heart, kidneys and intestine were aseptically removed.

*In vivo* and *ex vivo* luminescence images as well as fluorescence images of DiR-labeled LNPs were acquired using the IVIS Lumina XRMS Imaging System, following manufacturer’s instructions.

## Author Contributions

Conceptualization: AP, JH, JGW, JMO; Lipid molecular design and synthetic methodology: JH, AP, JGW; Lipid synthesis and characterization experiments: AP, JH, BB; LNP formulation and characterization: AT, TA, DDM; mRNA synthesis: DC; Biological and biochemical assays: AL, AGL, EM; Supervision: JGW, JMO, EPH; Writing the original draft: JH, AP, JGW; Manuscript review: all authors.

## Funding Sources

AP is funded by DIN2021-011799 financed by the Ministerio de Ciencia e Innovación MCIN/AEI / 10.13039/501100011033. This study was supported by Gobierno de Aragón (Spain) through project IDMF/2021/0009 (Nuevas tecnologías para el diseño y obtención de vacunas de ARN en Aragón).

## Notes

The authors declare the following competing financial interest(s): AP, JH, DDM, AT, JMO and JGW are inventors on patents related to this publication. All the authors declare to be employees of Certest Biotec.

## STATISTICAL ANALYSIS

Data are presented as mean ± SD or mean. Student’s t-test, one-way or two-way analysis of variance (ANOVA) followed by Tukey test was applied for comparison between two groups or among multiple groups using GraphPad Prism 10.0, respectively.

## ACKNOWLEDGEMENT

Authors would like to acknowledge the use of Servicios Científico Técnicos del CIBA (IACS-Universidad de Zaragoza) and Servicio de análisis bioquímicos de CIMA (Universidad de Navarra). Authors would like to extend their sincere gratitude to all the members of Certest Pharma group as well as Dr. Irene Maluenda-Borderas. Their valuable insights and expertise significantly contributed to the success of this research.

## Notes

### Summary of Updates

This version has been revised to improve the quality of some of the figures.

