## Supporting information for "Multicomponent Thiolactone-Based Ionizable Lipid Screening Platform for an Efficient and Tunable mRNA Delivery to the Lungs"

### Synthesis of ionizable lipids and their intermediates

#### General synthesis of intermediate (Bj).

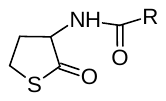

DL-homocysteine thiolactone hydrochloride (1.8 mmol) was dissolved in 5 mL of anhydrous dichloromethane at room temperature. Then triethylamine (1.8 mmol) was added followed by EDC hydrochloride (1.3 mmol), 4-(dimethylamino)pyridine (0.20 mmol) and carboxylic acid (1.0 mmol). The reaction mixture was stirred at room temperature overnight under an argon atmosphere. The organic layer was dried with anhydrous MgSO_4_, filtered, and evaporated under reduced pressure. The resulting residue was purified by flash chromatography (gradient of hexane/ethyl acetate: 100/0 to 0/100) to afford compound **B** as a pure product.

The pure compounds were characterized by mass spectroscopy.

**B1:** Theoretical [M+H]^+^= 384.29, experimental [M+H]^+^= 384.47

**B2:** Theoretical [M+H]^+^= 356.26, experimental [M+H]^+^= 356.44

**B3:** Theoretical [M+H]^+^=382.28, experimental [M+H]^+^=382.45

**B4:** Theoretical [M+H]^+^= 300.19, experimental [M+H]^+^= 300.32

**B5:** The synthesis of this compound requires one previous oxidation step to obtain the respective carboxylic acid. First, 2-octyl-1-dodecanol (1077 mg, 3.5 mmol) and periodic acid (4144 mg, 18 mmol) were dissolved in acetonitrile (28 ml). Then pyridinium chlorochromate (23 mg, 0.030 mmol) was added to the previous solution. The reaction mixture was stirred 16 hours at room temperature. Then, the solvent was evaporated under reduced pressure. Thus, the crude was dissolved in AcOEt and first washed twice with water and finally with brine. The organic phase was dried with anhydrous MgSO_4_, filtered and evaporated under reduced pressure. The residue was used in the next step without any further purification.

Theoretical [M+H]^+^= 412.32, experimental [M+H]^+^= 412.50

#### General synthesis of non-commercial acrylates (C_k_).

For those acrylates not available commercially the following general synthesis was followed:

Acryloyl chloride (4.0 mmol) was added dropwise to a cooled (0 ºC) solution of alcohol (4.4 mmol) and triethylamine (6.0 mmol) in anhydrous dichloromethane (15 mL). The resulting solution was protected from light and stirred overnight under an argon atmosphere, allowing it to warm to room temperature. The reaction mixture was followed to completion by TLC chromatography. Then, the reaction mixture was first washed twice with water and finally with brine. The organic phase was dried with anhydrous MgSO_4_, filtered and evaporated under reduced pressure. The resulting residue was purified by flash chromatography (gradient of hexane/ethyl acetate: 100/0 to 0/100) to afford the branched acrylate as a pure product.

**C1:** ^1^H NMR (400 MHz, CDCl_3_) δ: 6.39 (dd, J = 17.4, 1.5 Hz, 1H), 6.12 (dd, J = 17.3, 10.5 Hz, 1H), 5.81 (dd, J = 10.4, 1.5 Hz, 1H), 5.40 – 5.29 (m, 2H), 4.14 (t, J = 6.7 Hz, 2H), 2.01 (q, J = 6.5 Hz, 4H), 1.74 – 1.60 (m, 2H), 1.40 – 1.24 (m, 22H), 0.92 – 0.84 (m, 3H).

**C3:** ^1^H NMR (400 MHz, CDCl_3_) δ: 6.39 (dd, J = 17.4, 1.5 Hz, 1H), 6.12 (dd, J = 17.3, 10.4 Hz, 1H), 5.81 (dd, J = 10.4, 1.5 Hz, 1H), 4.06 (d, J = 5.8 Hz, 2H), 1.67 (t, J = 6.0 Hz, 1H), 1.36 – 1.24 (m, 16H), 0.94 – 0.84 (m, 6H)

**C8:** ^1^H NMR (400 MHz, CDCl_3_) δ: 6.38 (dd, J= 16.9 Hz J=1.6 Hz, 1H); 6.11 (dd, J=10.4 Hz J=16.9 Hz ,1H); 5.80 (dd, J=10.4 Hz J=1.6 Hz, 1H); 4.05 (d, J=5.6 Hz, 2H); 1.66 (m, 1H ); 1.27 (m, 24H); 0.87 (t, J=6.7 Hz, 3H)

**C9:** ^1^H NMR (400 MHz, CDCl_3_) δ: 6.39 (dd, J = 17.3, 1.5 Hz, 1H), 6.12 (dd, J = 17.3, 10.4 Hz, 1H), 5.81 (dd, J = 10.4, 1.5 Hz, 1H), 4.06 (d, J = 5.8 Hz, 2H), 1.58 (m, 1H), 1.36-1.23 (m, 30H), 0.92 – 0.84 (m, 6H).

#### Synthesis of lipid derivatives of A4B2C3 (Figure 3a)

The seven lipids were synthesized according to the general method describing in the article for the synthesis of STAAR lipids. Adding a final purification through flash chromatography with gradient elution from 100% of dichloromethane to 50% of 80/20/1 DCM/MeOH/ NH_4_OH (aq).

The pure compounds were characterized by mass spectroscopy.

**A4B2C3:** (74 mg, 78%). Theoretical [M+H]^+^=684.57, experimental [M+H]^+^=684.77.

**A4B2C8 (CP-LC-0729) (top-performing):** (84 mg**,** 76%). Written in the article.

**A4B2C9:** (85 mg**,** 71%). Theoretical [M+H]^+^=796.69, experimental [M+H]^+^=796.89.

**A4B4C3:** (74 mg, 77%). Theoretical [M+H]^+^=628.51, experimental [M+H]^+^=628.65.

**A4B4C8:** (75 mg, 73%). Theoretical [M+H]^+^=684.57, experimental [M+H]^+^=684.75.

**A4B4C9:** (90 mg, 81%). Theoretical [M+H]^+^=740.63, experimental [M+H]^+^=740.83.

**A4B5C3:** (85 mg, 77%). Theoretical [M+H]^+^=740.63, experimental [M+H]^+^=740.86.

**A4B5C8:** (89 mg, 75%). Theoretical [M+H]+=796.69, experimental [M+H]+=796.87.

**A4B5C9:** (95 mg, 74%) Theoretical [M+H]^+^=852.76, experimental [M+H]^+^=853.00.

#### Preparation of intermediates required for the synthesis of derivatives of CP-LC-0729 (01 to 05, Figure 3b)

##### Synthesis of intermediate 2-oxotetrahydrothiophene-3-carboxylic acid.

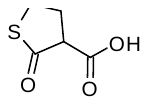

The product was synthesized according to the literature^1^.

^1^H NMR (400 MHz, CD_3_CN) δ: 2.55(dd, 1H), 3.50-3.46 (m, 2H), 2.43-2.61 (m, 2H).

##### Synthesis of intermediate 2-hexyldecyl 2-oxotetrahydrothiophene-3-carboxylate.

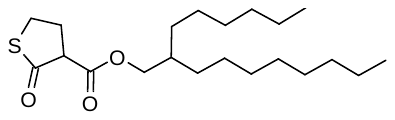

2-Oxotetrahydrothiophene-3-carboxylic acid (175 mg, 1.2 mmol) was dissolved in 5 mL of anhydrous dichloromethane at room temperature. Then 2-hexyl-1-decanol (299 mg, 1.0 mmol) was added followed by EDC hydrochloride (254 mg, 1.3 mmol) and 4-(dimethylamino)pyridine (25 mg, 0.2 mmol). The reaction mixture was stirred at room temperature overnight under an argon atmosphere. After that, the reaction mixture was first washed twice with water and finally with brine. The organic layer was dried with anhydrous MgSO_4_, filtered, and evaporated under reduced pressure. The resulting residue was purified by flash chromatography (gradient of hexane/dichloromethane: 100/0 to 0/100) to afford the product as a colorless liquid (271 mg, 73%).

^1^H NMR (400 MHz, CDCl_3_) δ: 4.15 – 4.00 (m, 2H), 3.57 – 3.45 (m, 2H), 3.36 (m, 1H), 2.64 (m, 1H), 2.50 (m, 1H), 1.70 – 1.62 (m, 1H), 1.27 (m, 24H), 0.92 – 0.84 (m, 6H).

##### Synthesis of intermediate N-(2-hexyldecyl)-2-oxotetrahydrothiophene-3-carboxamide.

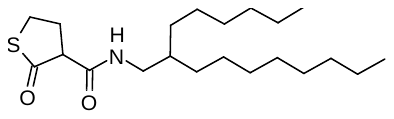

2-Oxotetrahydrothiophene-3-carboxylic acid (175 mg, 1.2 mmol) was dissolved in 5 mL of anhydrous dichloromethane at room temperature. Then 2-hexyldecan-1-amine (170 mg, 1.0 mmol) was added followed by EDC hydrochloride (254 mg, 1.3 mmol) and 4-(dimethylamino)pyridine (25 mg, 0.2 mmol). The reaction mixture was stirred at room temperature overnight under an argon atmosphere. After that, the reaction mixture was first washed twice with water and finally with brine. The organic layer was dried with anhydrous MgSO_4_, filtered, and evaporated under reduced pressure. The resulting residue was purified by flash chromatography (gradient of hexane/ethyl acetate: 100/0 to 0/100) to afford the product as a whitish solid (248 mg, 67%).

The pure compound was characterized by mass spectroscopy.

Theoretical [M+H]^+^= 370.28 , experimental [M+H]^+^= 370.46

##### 2-(Dimethylamino)ethyl (2-hexyldecanoyl)homocysteinate.

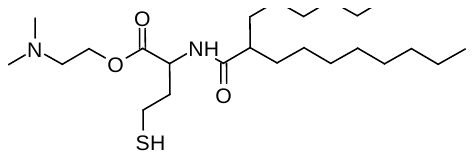

(2-Hexyldecanoyl)homocysteine thiolactone (**B2**) (160 mg ,0.36 mmol) was added over 2-(dimethylamino)ethanol (2.00 ml, 20 mmol), the reaction mixture was heated at 85ºC for 16 hours under argon atmosphere. Subsequently, the reaction mixture was first washed with 1M HCl, once with water and finally with brine. The organic layer was dried with anhydrous MgSO_4_, filtered, and evaporated under reduced pressure. The resultant residue was a mixture of B2 (45%) and the target product (55%). This residue was used in the next step without further purification.

The pure compound was characterized by mass spectroscopy. Theoretical [M+H]+=445.35, experimental [M+H]+=445.52

##### 1-(2-(Dimethylamino)ethyl) 3-(2-hexyldecyl) 2-(2-mercaptoethyl)malonate.

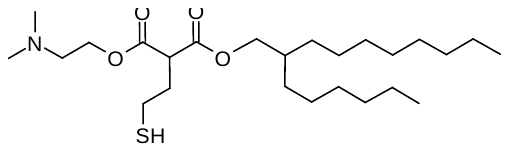

2-Hexyldecyl 2-oxotetrahydrothiophene-3-carboxylate (173 mg ,0.47 mmol) was added in 2-(dimethylamino)ethanol (2.59 ml, 26 mmol), the reaction mixture was heated at 85ºC during 16 hours under argon atmosphere. Subsequently, the reaction mixture was first washed with 1M HCl, once with water and finally with brine. The organic layer was dried with anhydrous MgSO_4_, filtered, and evaporated under reduced pressure. The resultant residue was a mixture of 1-(2-(dimethylamino)ethyl) 3-(2-hexyldecyl) 2-(2-mercaptoethyl)malonate (50%) and the target product (50%). This residue was used in the next step without further purification (91 mg, 43%).

The pure compound was characterized by mass spectroscopy. Theoretical [M+H]^+^=460.35, experimental [M+H]^+^=460.52

#### Synthesis lipid derivatives of CP-LC-0729 (01 to 05, Figure 3b).

Lipid CP-LC-0729 and three of its variants (CP-LC-0729-02, CP-LC-0729-03 and CP-LC-0729-04) were synthesized according to the general method described in the article for the synthesis of STAAR lipids using their corresponding initial products. Adding a final purification through flash chromatography with gradient elution from 100% of dichloromethane to 50% of 80/20/1 DCM/MeOH/ NH_4_OH (aq) allowed the obtention of these products in the following purities and yields:

**A4B2C8 (CP-LC-0729) (top-performing):** (84 mg**,** 76%). Written in the article.

Theoretical [M+H]^+^=740.63, experimental [M+H]^+^=740.84.

**CP-LC-0729-02:** A3, B2 and C8 were used as initial products. (85 mg, 75%).

Theoretical [M+H]^+^=754.65, experimental [M+H]^+^=754.80.

**CP-LC-0729-03:** A4, C8 and N-(2-hexyldecyl)-2-oxotetrahydrothiophene-3-carboxamide were used as initial products. (79 mg, 70%).

Theoretical [M+H]^+^=754.24, experimental [M+H]^+^=754.81.

**CP-LC-0729-04:** A4, C8 and 2-hexyldecyl 2-oxotetrahydrothiophene-3-carboxylate were used as initial products. (69 mg, 61%).

Theoretical [M+H]^+^=755.63, experimental [M+H]^+^=755.83.

The other two lipids required of alternative synthetic methods as follows:

**CP-LC-0729-01:** Intermediate 2-(dimethylamino)ethyl (2-hexyldecanoyl)homocysteinate (89 mg, 0.2 mmol) (described in the previous section) and acrylate C8 (89 mg, 0.3 mmol) were dissolved in 0.4 ml tetrahydrofuran. Then, triethylamine was added (31 mg, 0.3 mmol). The reaction mixture was stirred at room temperature for 16 hours under argon atmosphere. Later, the crude was purified through flash chromatography with gradient elution from 100% of dichloromethane to 50% of 80/20/1 DCM/MeOH/ NH_4_OH (aq). (62 mg, 42%)

Theoretical [M+H]^+^=741.62, experimental [M+H]^+^=741.84.

**CP-LC-0729-05:** Intermediate 1-(2-(dimethylamino)ethyl) 3-(2-hexyldecyl) 2-(2-mercaptoethyl)malonate (91 mg, 0.2 mmol) (described in the previous section) and C8 (119mg, 0.4 mmol) were dissolved in 0.4 ml tetrahydrofuran. Then, triethylamine was added (56 mg, 0.4 mmol). The reaction mixture was stirring at room temperature during 16 hours under argon atmosphere. Later, the crude was purified through flash chromatography with gradient elution from 100% of dichloromethane to 50% of 80/20/1 DCM/MeOH/ NH_4_OH (aq). (56 mg, 37%)

Theoretical [M+H]^+^=756.62, experimental [M+H]^+^=756.78.

### NMR spectra

All NMR spectra were obtained using a Bruker Avance 400 MHz device and deuterated solvents (CDCl_3_, Sigma Aldrich; DMSO-d_6_, Sigma Aldrich)

### SUPPORTING FIGURES AND TABLES

GpppAGGAGGCACAGACACCAAGGACAGAGACGCΨGGCΨAGGCCGCCCΨCCCCACΨGΨΨACCAACGCCACCAΨGGAGGACGCCAAGAACAΨCAAGAAGGGCCCCGCCCCCΨΨCΨACCCCCΨGGAGGACGGCACCGCCGGCGAGCAGCΨGCACAAGGCCAΨGAAGAGGΨACGCCCΨGGΨGCCCGGCACCAΨCGCCΨΨCACCGACGCCCACAΨCGAGGΨGGACAΨCACCΨACGCCGAGΨACΨΨCGAGAΨGAGCGΨGAGGCΨGGCCGAGGCCAΨGAAGAGGΨACGGCCΨGAACACCAACCACAGGAΨCGΨGGΨGΨGCAGCGAGAACAGCCΨGCAGΨΨCΨΨCAΨGCCCGΨGCΨGGGCGCCCΨGΨΨCAΨCGGCGΨGGCCGΨGGCCCCCGCCAACGACAΨCΨACAACGAGAGGGAGCΨGCΨGAACAGCAΨGGGCAΨCAGCCAGCCCACCGΨGGΨGΨΨCGΨGAGCAAGAAGGGCCΨGCAGAAGAΨCCΨGAACGΨGCAGAAGAAGCΨGCCCAΨCAΨCCAGAAGAΨCAΨCAΨCAΨGGACAGCAAGACCGACΨACCAGGGCΨΨCCAGAGCAΨGΨACACCΨΨCGΨGACCAGCCACCΨGCCCCCCGGCΨΨCAACGAGΨACGACΨΨCGΨGCCCGAGAGCΨΨCGACAGGGACAAGACCAΨCGCCCΨGAΨCAΨGAACAGCAGCGGCAGCACCGGCCΨGCCCAAGGGCGΨGGCCCΨGCCCCACAGGACCGCCΨGCGΨGAGGΨΨCAGCCACGCCAGGGACCCCAΨCΨΨCGGCAACCAGAΨCAΨCCCCGACACCGCCAΨCCΨGAGCGΨGGΨGCCCΨΨCCACCACGGCΨΨCGGCAΨGΨΨCACCACCCΨGGGCΨACCΨGAΨCΨGCGGCΨΨCAGGGΨGGΨGCΨGAΨGΨACAGGΨΨCGAGGAGGAGCΨGΨΨCCΨGAGGAGCCΨGCAGGACΨACAAGAΨCCAGAGCGCCCΨGCΨGGΨGCCCACCCΨGΨΨCAGCΨΨCΨΨCGCCAAGAGCACCCΨGAΨCGACAAGΨACGACCΨGAGCAACCΨGCACGAGAΨCGCCAGCGGCGGCGCCCCCCΨGAGCAAGGAGGΨGGGCGAGGCCGΨGGCCAAGAGGΨΨCCACCΨGCCCGGCAΨCAGGCAGGGCΨACGGCCΨGACCGAGACCACCAGCGCCAΨCCΨGAΨCACCCCCGAGGGCGACGACAAGCCCGGCGCCGΨGGGCAAGGΨGGΨGCCCΨΨCΨΨCGAGGCCAAGGΨGGΨGGACCΨGGACACCGGCAAGACCCΨGGGCGΨGAACCAGAGGGGCGAGCΨGΨGCGΨGAGGGGCCCCAΨGAΨCAΨGAGCGGCΨACGΨGAACAACCCCGAGGCCACCAACGCCCΨGAΨCGACAAGGACGGCΨGGCΨGCACAGCGGCGACAΨCGCCΨACΨGGGACGAGGACGAGCACΨΨCΨΨCAΨCGΨGGACAGGCΨGAAGAGΨCΨGAΨCAAGΨACAAGGGCΨACCAGGΨGGCCCCCGCCGAGCΨGGAGAGCAΨCCΨGCΨGCAGCACCCCAACAΨCΨΨCGACGCCGGCGΨGGCCGGCCΨGCCCGACGACGACGCCGGCGAGCΨGCCCGCCGCCGΨGGΨGGΨGCΨGGAGCACGGCAAGACCAΨGACCGAGAAGGAGAΨCGΨGGACΨACGΨGGCCAGCCAGGΨGACCACCGCCAAGAAGCΨGAGGGGCGGCGΨGGΨGΨΨCGΨGGACGAGGΨGCCCAAGGGCCΨGACCGGCAAGCΨGGACGCCAGGAAGAΨCAGGGAGAΨCCΨGAΨCAAGGCCAAGAAGGGCGGCAAGAΨCGCCGΨGΨGAΨGAGCΨCGCΨΨΨCΨΨGCΨGΨCCAAΨΨΨCΨAΨΨAAAGGΨΨCCΨΨΨGΨΨCCCΨAAGΨCCAACΨACΨAAACΨGGGGGAΨAΨΨAΨGAAGGGCCΨΨGAGCAΨCΨGGAΨΨCΨGCCΨAAΨAAAAAACAΨΨΨAΨΨΨΨCAΨΨGCGCΨCGCΨΨΨCΨΨGCΨGΨCCAAΨΨΨCΨAΨΨAAAGGΨΨCCΨΨΨGΨΨCCCΨAAGΨCCAACΨACΨAAACΨGGGGGAΨAΨΨAΨGAAGGGCCΨΨGAGCAΨCΨGGAΨΨCΨGCCΨAAΨAAAAAACAΨΨΨAΨΨΨΨCAΨΨGCAAAAAAAAAAAAAAAAAAAAAAAAAAAAAAGAAAAAAAAAAAAAAAAAAAAAAAAAAAAAAAAAAAAAAAAAAAAAAAAAAAAAAAAAAAAAAAAAAAAAA

**Figure S1.** Firefly luciferase mRNA complete sequence. Where Ψ= 1-methyl-3'-pseudouridylyl^2,3^.

^1^H NMR (400 MHz, CDCl_3_) CP-LC-0729 (A4B2C8)

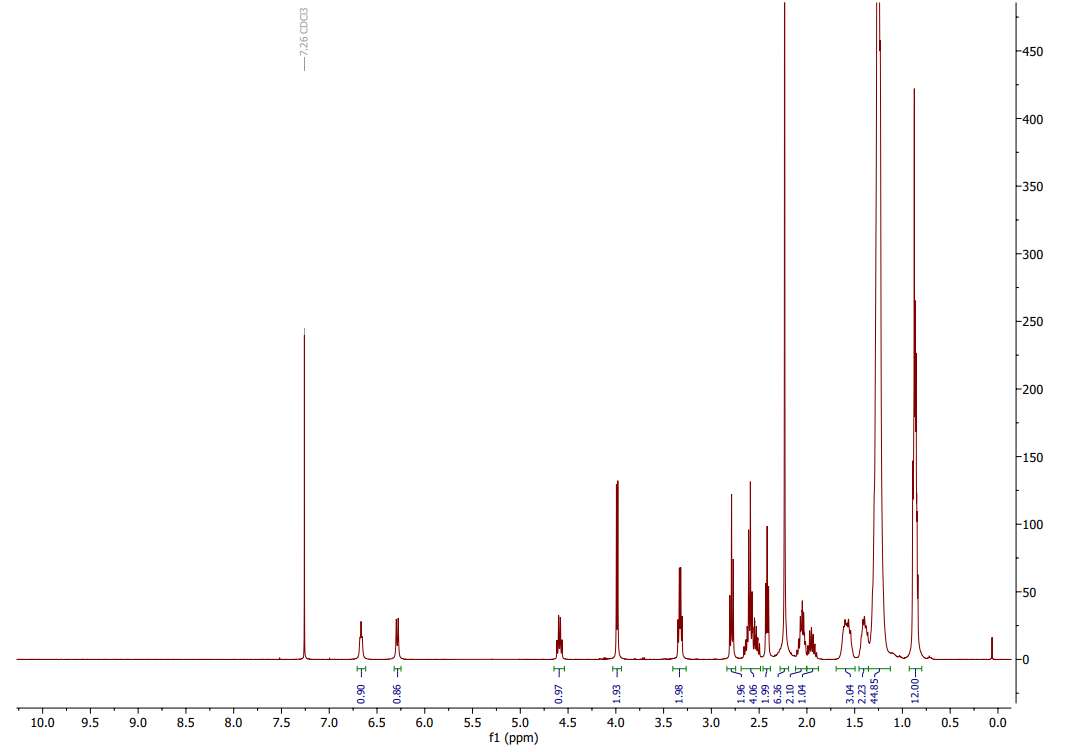
Mass Spectroscopy Spectrum CP-LC-0729 (A4B2C8)

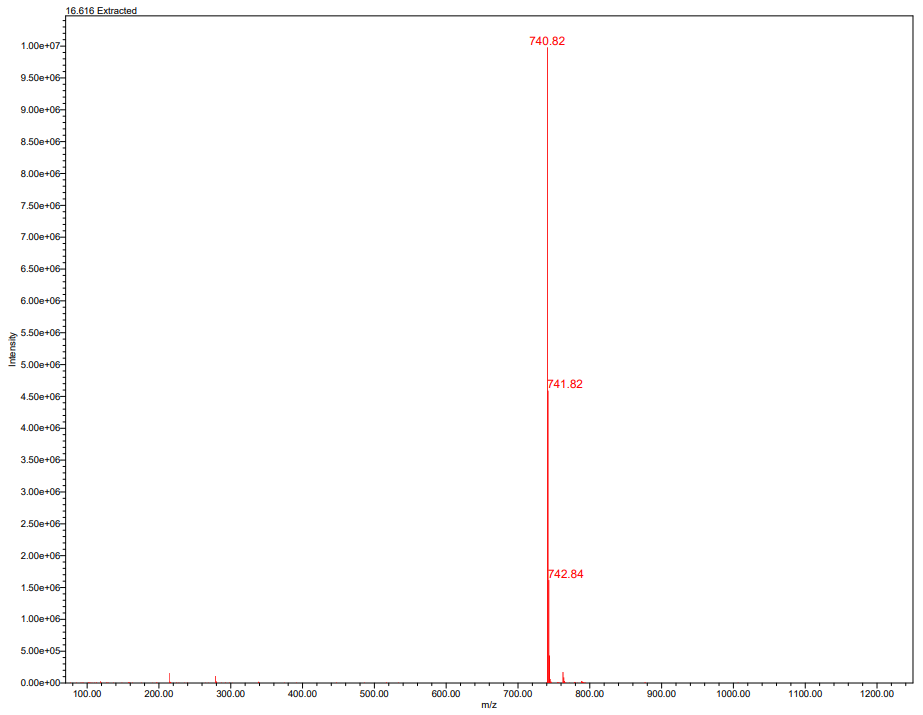

^1^H NMR (400 MHz, DMSO-d_6_) (+) CP-LC-0729 )

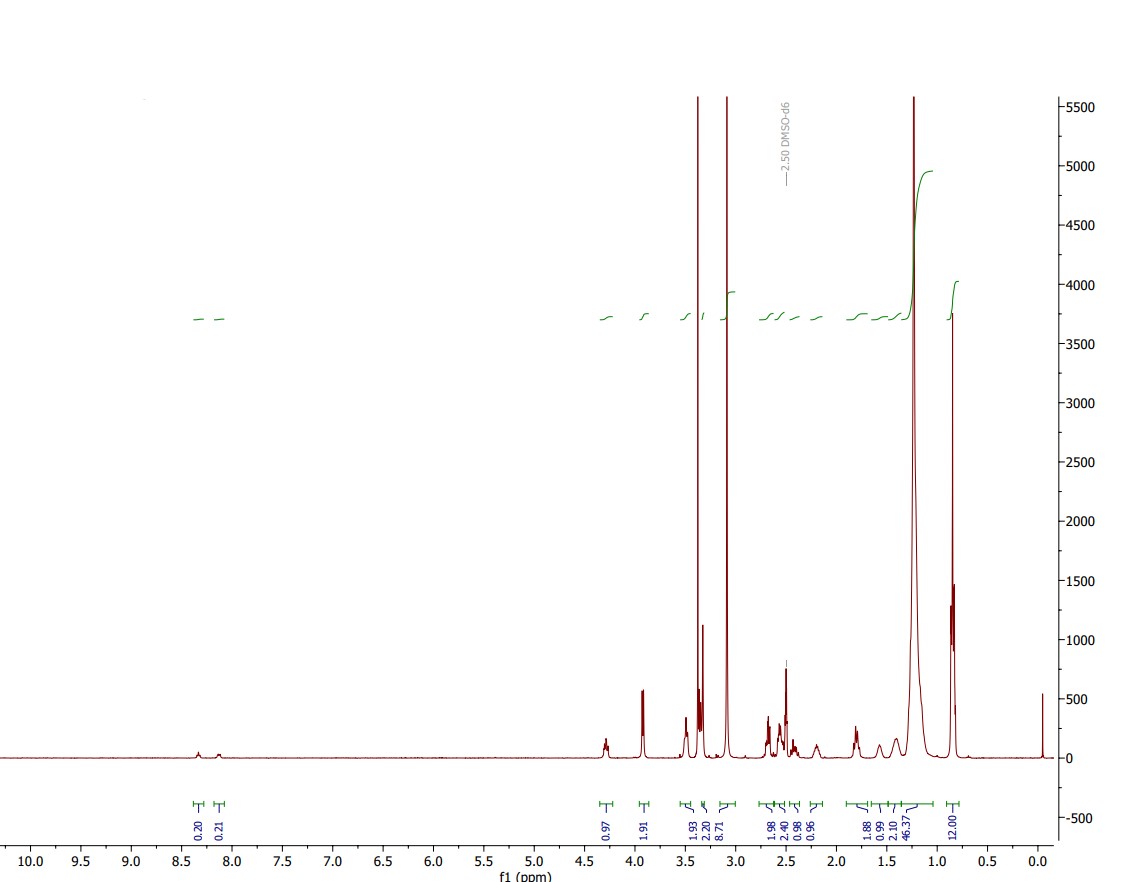

Mass Spectroscopy Spectrum CP-LC-0729 (A4B2C8)

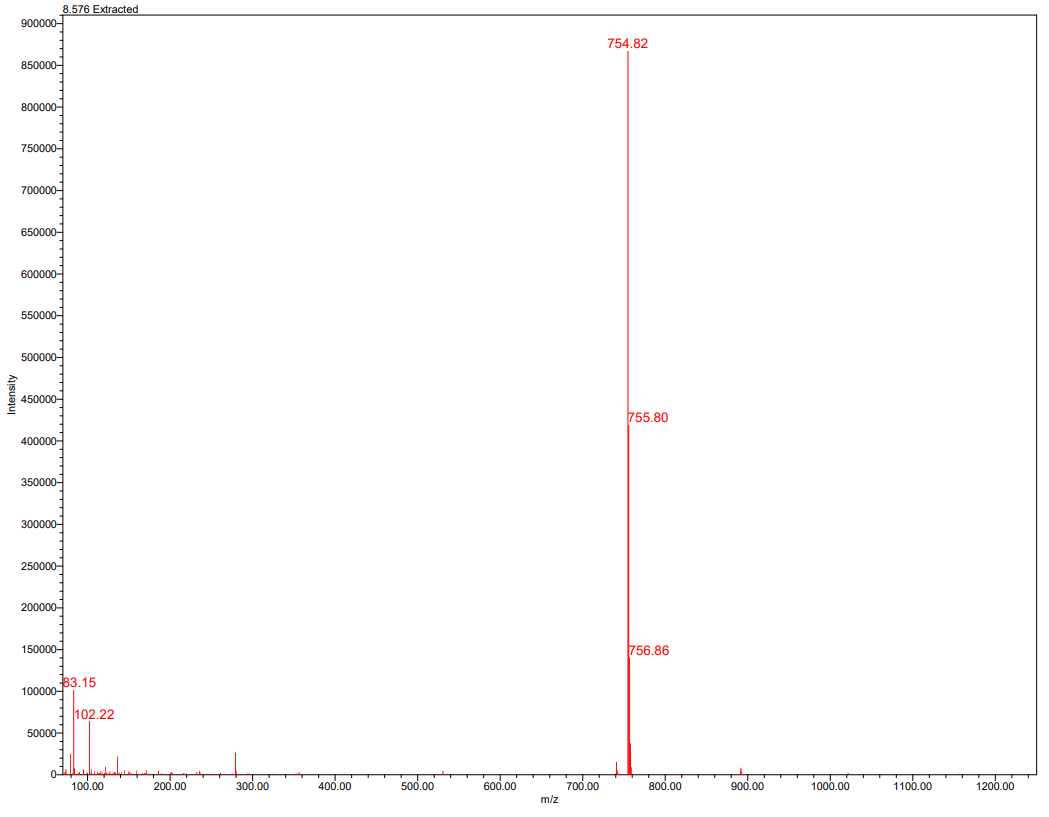

**Figure S2**. Mass spectrum and ^1^H NMR of purified CP-LC-0729 and (+) CP-LC-0729.

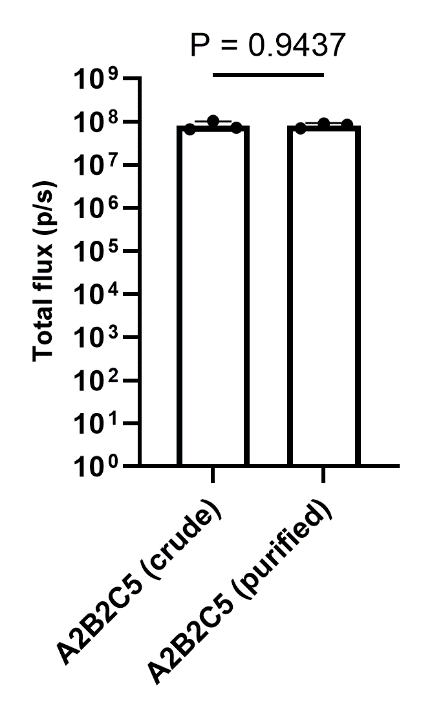

**Figure S3. Comparison of purified and crude LNPs of lipid A2B2C5 (in vivo expression).** In vivo mLuc expression (n = 3 biologically independent samples). Data are presented as mean values ± SD. Mice were i.m. injected with mLuc-loaded LNPs at an mRNA dose of 0.05 mg/kg. Luminescence imaging was performed at 4 h post-treatment and total flux was quantified. Student’s t-test was used.

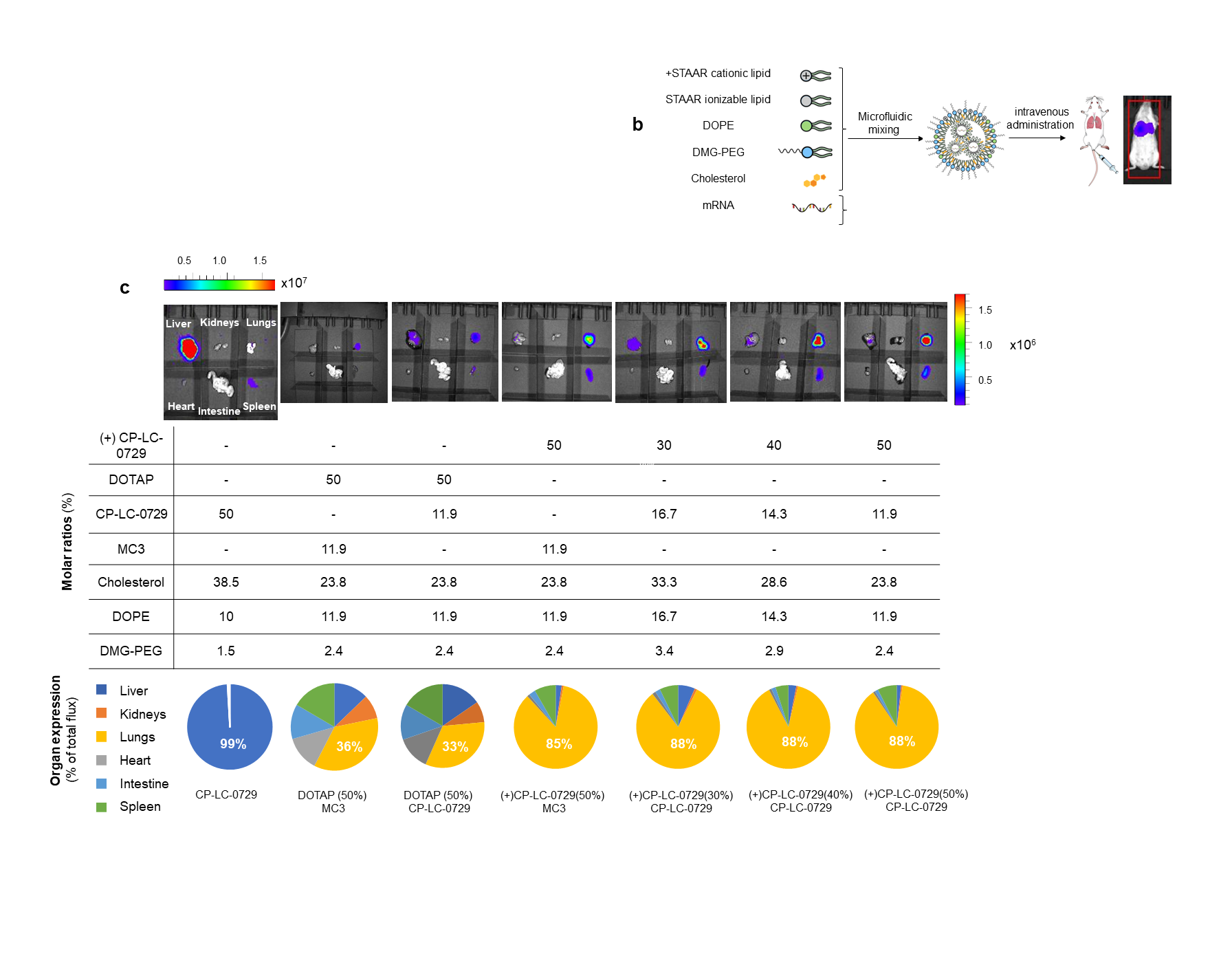
**Figure S4.** *Ex vivo* luminescence of major organs from mice and their quantification (n= 3 biologically independent samples). Mice were i.v. injected with mRNA-Luc-loaded LNPs at an mRNA dose of 0.2 mg/kg. Images were acquired at 4h post-treatment. Below the images, a table with the formulation details of LNPs and pie charts that show a profile of the tissue-specificity of mLuc expression in the lungs, heart, liver, intestine, spleen and kidneys and the quantification of the majority percent expression detected in each LNP.

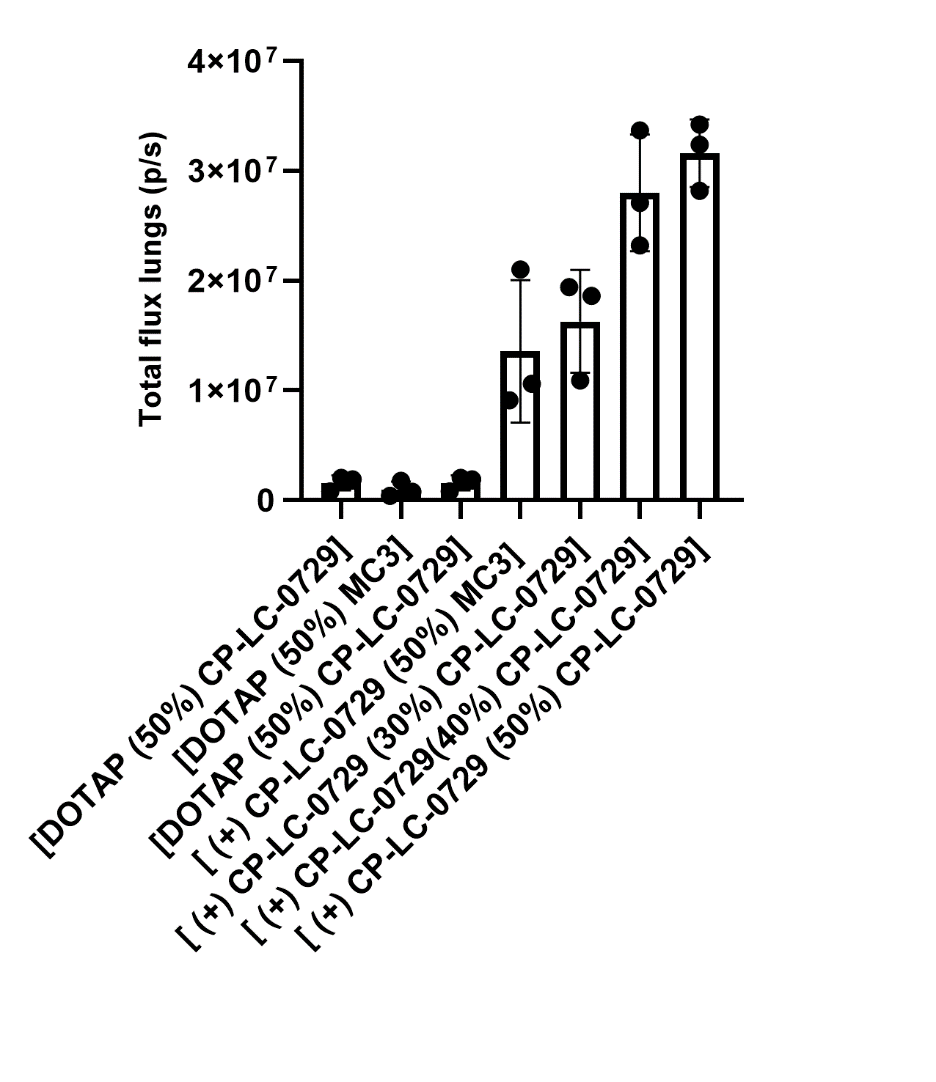

**Figure S5.** Quantification of the total luminescence flux produced by the LNPs containing cationic lipids in the lungs (n= 3 biologically independent samples). Mice were i.v injected with mRNA-Luc-loaded LNPs at an mRNA dose of 0.2 mg/kg. Images and their quantification were taken at 4h post-treatment. Data are presented as mean values ± SD. One-way ANOVA with Tukey’s correction was used.

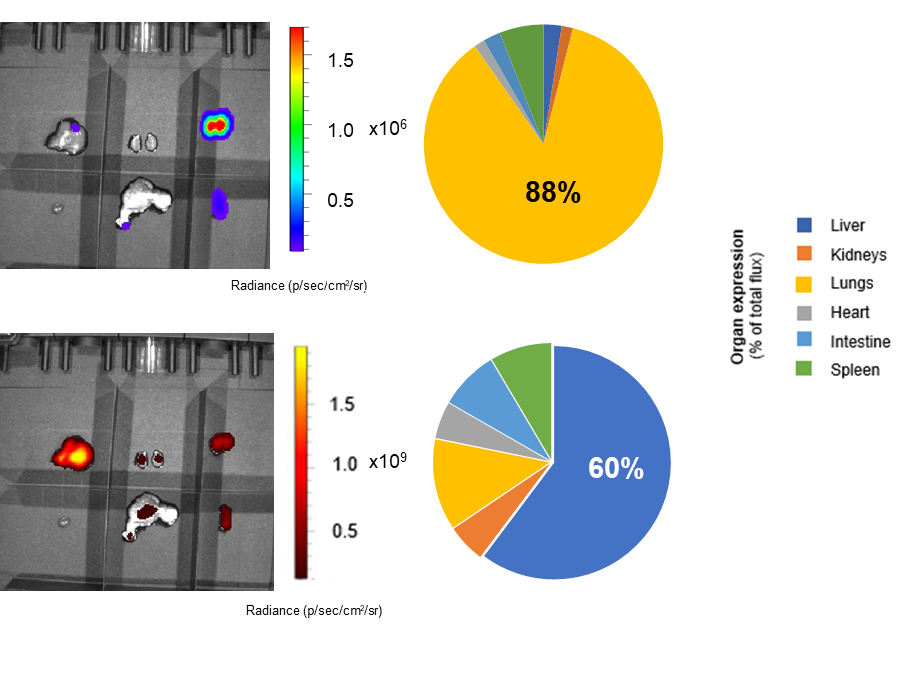

**Figure S6.** Images of the luminescence (above) and fluorescence (below) produced by [(+) CP-LC-0729 (50%) CP-LC-0729]-DiR LNP in the liver, kidneys, lungs, heart, intestine and spleen (n= 3 biologically independent samples). Mice were i.v. injected with mLuc-loaded LNPs at an mRNA dose of 0.2 mg/kg. Images were taken at 4h post-treatment. Beside the images, pie charts profile the tissue-specificity of mLuc (above) and fluorescence (below) expression in the lungs, heart, liver, intestine, spleen and kidneys and quantify the majority percent expression detected.

**Table S1.** A compilation of the detected m/z values in the mass spectra of STAAR Lipids during two optimization phases of screening:

|  |  | Theoretical [M+H]^+^ | Experimental  [M+H]^+^ |
| --- | --- | --- | --- |
| Phase 1 optimization | A1B1C1 | 808.69 | 808.84 |
|  | A1B1C2 | 670.55 | 670.73 |
|  | A1B1C3 | 726.62 | 726.81 |
|  | A1B1C4 | 754.64 | 754.83 |
|  | A1B1C5 | 726.61 | 726.84 |
|  | A1B1C6 | 670.55 | 670.72 |
|  | A1B1C7 | 670.55 | 670.76 |
|  | A1B2C1 | 780.66 | 780.80 |
|  | A1B2C2 | 642.52 | 642.68 |
|  | A1B2C3 | 698.60 | 698.76 |
|  | A1B2C4 | 726.61 | 726.82 |
|  | A1B2C5 | 698.58 | 698.82 |
|  | A1B2C6 | 642.52 | 642.67 |
|  | A1B2C7 | 642.52 | 642.69 |
|  | A1B3C1 | 806.68 | 806.82 |
|  | A1B3C2 | 668.53 | 668.73 |
|  | A1B3C3 | 724.60 | 724.80 |
|  | A1B3C4 | 752.63 | 752.87 |
|  | A1B3C5 | 724.60 | 724.85 |
|  | A1B3C6 | 668.53 | 668.73 |
|  | A1B3C7 | 668.53 | 668.73 |
| Phase 2  optimization | A2B2C1 | 794.68 | 794.87 |
|  | A2B2C2 | 656.53 | 656.69 |
|  | A2B2C3 | 712.60 | 712.79 |
|  | A2B2C4 | 740.63 | 740.82 |
|  | A2B2C5 | 712.60 | 712.79 |
|  | A2B2C6 | 656.53 | 656.70 |
|  | A2B2C7 | 656.53 | 656.70 |
|  | A3B2C1 | 780.66 | 780.82 |
|  | A3B2C2 | 642.52 | 642.64 |
|  | A3B2C3 | 698.59 | 698.80 |
|  | A3B2C4 | 726.61 | 726.82 |
|  | A3B2C5 | 698.58 | 698.78 |
|  | A3B2C6 | 642.03 | 642.67 |
|  | A3B2C7 | 642.52 | 642.67 |
|  | A4B2C1 | 766.25 | 766.81 |
|  | A4B2C2 | 628.50 | 628.66 |
|  | A4B2C3 | 684.57 | 684.75 |
|  | A4B2C4 | 712.60 | 712.74 |
|  | A4B2C5 | 684.57 | 684.74 |
|  | A4B2C6 | 628.50 | 628.65 |
|  | A4B2C7 | 628.50 | 628.65 |
|  | A5B2C1 | 821.69 | 821.84 |
|  | A5B2C2 | 683.55 | 683.70 |
|  | A5B2C3 | 739.61 | 739.81 |
|  | A5B2C4 | 767.64 | 767.81 |
|  | A5B2C5 | 739.61 | 739.79 |
|  | A5B2C6 | 683.55 | 683.71 |
|  | A5B2C7 | 683.55 | 683.70 |
|  | A6B2C1 | 777.62 | 777.77 |
|  | A6B2C2 | 665.50 | 665.71 |
|  | A6B2C3 | 721.57 | 721.77 |
|  | A6B2C4 | 749.59 | 749.86 |
|  | A6B2C5 | 721.56 | 721.79 |
|  | A6B2C6 | 665.50 | 665.69 |
|  | A6B2C7 | 665.50 | 665.71 |
|  | A7B2C1 | 822.31 | 822.86 |
|  | A7B2C2 | 684.53 | 684.72 |
|  | A7B2C3 | 740.60 | 740.79 |
|  | A7B2C4 | 768.62 | 768.80 |
|  | A7B2C5 | 740.59 | 740.80 |
|  | A7B2C6 | 684.53 | 684.70 |
|  | A7B2C7 | 684.53 | 684.68 |
|  | A8B2C1 | 792.66 | 792.78 |
|  | A8B2C2 | 654.52 | 654.68 |
|  | A8B2C3 | 710.59 | 710.77 |
|  | A8B2C4 | 738.61 | 738.83 |
|  | A8B2C5 | 710.58 | 710.77 |
|  | A8B2C6 | 654.52 | 654.69 |
|  | A8B2C7 | 654.52 | 654.69 |
|  | A9B2C1 | 806.68 | 806.84 |
|  | A9B2C2 | 668.53 | 668.73 |
|  | A9B2C3 | 724.60 | 724.78 |
|  | A9B2C4 | 752.63 | 752.85 |
|  | A9B2C5 | 724.60 | 724.83 |
|  | A9B2C6 | 668.53 | 668.74 |
|  | A9B2C7 | 668.53 | 668.73 |
|  | A10B2C1 | 840.69 | 840.86 |
|  | A10B2C2 | 702.54 | 702.74 |
|  | A10B2C3 | 758.60 | 758.79 |
|  | A10B2C4 | 786.64 | 786.83 |
|  | A10B2C5 | 758.61 | 758.80 |
|  | A10B2C6 | 702.54 | 702.73 |
|  | A10B2C7 | 702.54 | 702.75 |
|  | A11B2C1 | 806.68 | 806.85 |
|  | A11B2C2 | 668.54 | 668.73 |
|  | A11B2C3 | 724.60 | 724.81 |
|  | A11B2C4 | 752.63 | 752.83 |
|  | A11B2C5 | 724.60 | 724.82 |
|  | A11B2C6 | 668.54 | 668.71 |
|  | A11B2C7 | 668.54 | 668.72 |

**Table S2.** Characterization of formulated LNPs in screening phases 1 and 2.

|  |  | Particle  size (nm) | P.D.I. | Zeta potential (mV) | %EE |
| --- | --- | --- | --- | --- | --- |
| Phase 1 screening | A1B1C1 | 221.2 | 0.26 | 9.1 | 98.4 |
|  | A1B1C2 | 184.5 | 0.18 | 8.9 | 99.7 |
|  | A1B1C3 | 185.2 | 0.28 | 6.9 | 96.5 |
|  | A1B1C4 | 232.3 | 0.32 | 8.1 | 98.6 |
|  | A1B1C5 | 247.8 | 0.24 | 8.3 | 98.3 |
|  | A1B1C6 | 183.2 | 0.19 | 8.0 | 99.7 |
|  | A1B1C7 | 177.3 | 0.18 | 9.4 | 99.8 |
|  | A1B2C1 | 262.1 | 0.24 | 4.8 | 97.4 |
|  | A1B2C2 | 189.2 | 0.14 | -6.1 | 92.3 |
|  | A1B2C3 | 142.9 | 0.11 | -6.8 | 89.5 |
|  | A1B2C4 | 271.4 | 0.25 | 4.1 | 98.6 |
|  | A1B2C5 | 208.4 | 0.16 | 0.6 | 98.9 |
|  | A1B2C6 | 215.1 | 0.19 | -1.4 | 98.7 |
|  | A1B2C7 | 209.9 | 0.16 | 0.7 | 97.3 |
|  | A1B3C1 | 297.3 | 0.36 | 8.5 | 97.4 |
|  | A1B3C2 | 190.7 | 0.21 | 11.7 | 99.6 |
|  | A1B3C3 | 175.6 | 0.21 | 5.1 | 98.9 |
|  | A1B3C4 | 264.2 | 0.32 | 11.0 | 98.7 |
|  | A1B3C5 | 240.7 | 0.23 | 13.7 | 98.7 |
|  | A1B3C6 | 206.4 | 0.18 | 13.4 | 99.4 |
|  | A1B3C7 | 204.0 | 0.19 | 12.5 | 99.7 |
| Phase 2 screening | A2B2C1 | 143.8 | 0.13 | -5.9 | 60.6 |
|  | A2B2C2 | 148.5 | 0.23 | -9.1 | 35.5 |
|  | A2B2C3 | 145.1 | 0.10 | -9.4 | 50.0 |
|  | A2B2C4 | 156.2 | 0.17 | -7.0 | 65.2 |
|  | A2B2C5 | 146.7 | 0.15 | -9.6 | 57.4 |
|  | A2B2C6 | 161.6 | 0.21 | -5.9 | 64.0 |
|  | A2B2C7 | 136.9 | 0.16 | -6.1 | 56.2 |
|  | A3B2C1 | 193 | 0.12 | -12.3 | 76.3 |
|  | A3B2C2 | 207.1 | 0.14 | -11.1 | 49.7 |
|  | A3B2C3 | 165 | 0.25 | -13.4 | 50.0 |
|  | A3B2C4 | 191.9 | 0.15 | -11.5 | 75.1 |
|  | A3B2C5 | 186.3 | 0.14 | -7.5 | 80.1 |
|  | A3B2C6 | 143.8 | 0.17 | -16.3 | 57.8 |
|  | A3B2C7 | 203.8 | 0.16 | -10.2 | 65.1 |
|  | A4B2C1 | 225.8 | 0.17 | -1.4 | 73.2 |
|  | A4B2C2 | 171.9 | 0.13 | -3.0 | 67.4 |
|  | A4B2C3 | 109.1 | 0.15 | -8.5 | 95.9 |
|  | A4B2C4 | 201.9 | 0.19 | -1.7 | 65.2 |
|  | A4B2C5 | 204.3 | 0.17 | -0.5 | 44.4 |
|  | A4B2C6 | 197.0 | 0.11 | -0.4 | 80.1 |
|  | A4B2C7 | 181.7 | 0.12 | -2.2 | 68.1 |
|  | A5B2C1 | 231.7 | 0.31 | 4.9 | 98.7 |
|  | A5B2C2 | 206.7 | 0.35 | -6.5 | 85.5 |
|  | A5B2C3 | 148.2 | 0.14 | -7.2 | 81.3 |
|  | A5B2C4 | 186.8 | 0.15 | -1.6 | 92.4 |
|  | A5B2C5 | 217.1 | 0.22 | -2.7 | 91.2 |
|  | A5B2C6 | 135.2 | 0.11 | 4.8 | 87.0 |
|  | A5B2C7 | 158.7 | 0.18 | 3.4 | 86.7 |
|  | A6B2C1 | 211.3 | 0.10 | -7.4 | 47.1 |
|  | A6B2C2 | 138.8 | 0.14 | -8.5 | 41.9 |
|  | A6B2C3 | 132.0 | 0.14 | -10.9 | 44.3 |
|  | A6B2C4 | 187.3 | 0.13 | -12.6 | 58.1 |
|  | A6B2C5 | 161.7 | 0.18 | -17.2 | 44.3 |
|  | A6B2C6 | 176.7 | 0.15 | -10.3 | 36.6 |
|  | A6B2C7 | 147.0 | 0.15 | -8.3 | 39.0 |
|  | A7B2C1 | 169.3 | 0.18 | -12.2 | 41.0 |
|  | A7B2C2 | 164.8 | 0.11 | -24.6 | 42.5 |
|  | A7B2C3 | 190.3 | 0.27 | -10.6 | 51.0 |
|  | A7B2C4 | 198.1 | 0.16 | -23.7 | 58.8 |
|  | A7B2C5 | 200.6 | 0.22 | -22.3 | 34.3 |
|  | A7B2C6 | 204.1 | 0.17 | -9.7 | 44.9 |
|  | A7B2C7 | 228.8 | 0.24 | -23.1 | 53.7 |
|  | A8B2C1 | 216.2 | 0.23 | 1.7 | 84.5 |
|  | A8B2C2 | 179.0 | 0.27 | -20.9 | 83.3 |
|  | A8B2C3 | 135.9 | 0.12 | -11.1 | 69.8 |
|  | A8B2C4 | 174.1 | 0.11 | -0.9 | 87.2 |
|  | A8B2C5 | 221.3 | 0.19 | 4.5 | 64.6 |
|  | A8B2C6 | 196.7 | 0.13 | -7.0 | 92.8 |
|  | A8B2C7 | 156.7 | 0.15 | -2.4 | 72.8 |
|  | A9B2C1 | 146.9 | 0.13 | 1.0 | 93.4 |
|  | A9B2C2 | 175.0 | 0.16 | 1.1 | 53.6 |
|  | A9B2C3 | 151.2 | 0.12 | -1.0 | 92.3 |
|  | A9B2C4 | 192.7 | 0.16 | 10.0 | 92.7 |
|  | A9B2C5 | 177.3 | 0.16 | 5.9 | 94.6 |
|  | A9B2C6 | 168.7 | 0.20 | 7.0 | 80.0 |
|  | A9B2C7 | 142.0 | 0.21 | 3.7 | 63.7 |
|  | A10B2C1 | 184.8 | 0.16 | 2.5 | 97.5 |
|  | A10B2C2 | 153.3 | 0.20 | -8.2 | 87.1 |
|  | A10B2C3 | 158.4 | 0.12 | -9.3 | 71.7 |
|  | A10B2C4 | 194.1 | 0.16 | -1.6 | 96.0 |
|  | A10B2C5 | 227.4 | 0.29 | -2.9 | 90.2 |
|  | A10B2C6 | 185.7 | 0.20 | -0.8 | 93.6 |
|  | A10B2C7 | 171.7 | 0.14 | 0.8 | 89.6 |
|  | A11B2C1 | 142.7 | 0.12 | 0.3 | 90.8 |
|  | A11B2C2 | 156.1 | 0.11 | -11.2 | 43.9 |
|  | A11B2C3 | 229.1 | 0.20 | -1.4 | 72.0 |
|  | A11B2C4 | 163.6 | 0.13 | 1.2 | 87.6 |
|  | A11B2C5 | 193.9 | 0.21 | -2.1 | 87.5 |
|  | A11B2C6 | 223.3 | 0.27 | -11.1 | 74.8 |
|  | A11B2C7 | 251.3 | 0.32 | -14.0 | 47.3 |

LNPs were formulated by pipetting mixing. Aqueous and organic solutions were mixed by pipetting the aqueous solution into the organic solution by vigorous pipetting for about 10 seconds. The resulting lipid particles encapsulating mRNAs were immediately neutralized with a Tris buffered solution with 15% sucrose. Lipid particles were characterized by measuring their size distribution, polydispersity and Z-potential by dynamic light scattering (DLS). mRNA encapsulation was assessed by Quant-IT® Ribogreen following the manufacturer’s instructions.

**Table S3.** LNP formulation controls used.

|  | Components | Molar Ratio | N/P ratio (mol/mol) |
| --- | --- | --- | --- |
| MC3 Benchmark | MC3/DSPC/Chol/DMG-PEG | 50:10:38.5:1.5 | 6 |
| MC3 (DOPE) | MC3/DOPE/Chol/DMG-PEG | 50:10:38.5:1.5 | 6 |

**Table S4.** Characterization of formulated LNPs by microfluidics after screening phases.

|  | Particle  size (nm) | P.D.I. | Zeta potential (mV) | Apparent  pKa | %EE |
| --- | --- | --- | --- | --- | --- |
| A4B4C3 | 133.3 | 0.18 | -5.9 | 6.72 | 90.2 |
| A4B4C8 | 127.4 | 0.09 | -6.6 | 6.96 | 97.4 |
| A4B4C9 | 104.8 | 0.15 | -5.7 | 7.01 | 96.2 |
| A4B2C3 | 109.1 | 0.15 | -8.4 | 6.82 | 96.0 |
| **A4B2C8 (CP-LC-0729)** | **91.5** | **0.15** | **-7.7** | **6.78** | **95.1** |
| A4B2C9 | 86.12 | 0.12 | -9.1 | 6.89 | 94.1 |
| A4B5C3 | 107.8 | 0.20 | -5.3 | 6.76 | 97.5 |
| A4B5C8 | 98.47 | 0.26 | -10.4 | 6.63 | 96.5 |
| A4B5C9 | 90.29 | 0.14 | -7.8 | 6.61 | 94.4 |
| MC3 (DOPE) | 124.6 | 0.22 | -11.0 | 6.40 | 92.0 |
| MC3 (DSPC) | 86.53 | 0.28 | -13.0 | 6.15 | 96.3 |
| CP-LC-0729-01 | 72.7 | 0.12 | -10.3 | 6.12 | 93.8 |
| CP-LC-0729-02 | 90.6 | 0.09 | -8.2 | 6.81 | 96.1 |
| CP-LC-0729-03 | 76.3 | 0.11 | -5.9 | 7.06 | 97.1 |
| CP-LC-0729-04 | 87.7 | 0.15 | -9.5 | 7.18 | 98.0 |
| CP-LC-0729-05 | 80.6 | 0.25 | -18.0 | 5.48 | 92.9 |

LNPs were formulated by microfluidic mixing. The resulting lipid particles encapsulating mRNAs were dialyzed overnight against a pH 8 Tris buffer solution containing 15% sucrose. Lipid particles were characterized by measuring their size distribution, polydispersity and Z-potential by dynamic light scattering (DLS). mRNA encapsulation was assessed by Quant-IT® Ribogreen following the manufacturer’s instructions. Apparent pKa was calculated using a 6-(p-toluidinyl)naphthalene-2-sulfonic acid (TNS) assay.

**Table S5.** LNP formulation used in lung targeting strategy by microfluidics.

|  | Components | Molar Ratio | N/P ratio (w/w) |
| --- | --- | --- | --- |
| [DOTAP (50%) CP-LC-0729] | DOTAP/ CP-LC-0729 /DOPE/Chol/DMG-PEG/DiR | 50/11.9/11.9/23.8/2.4/0 | 10 |
| [ (+) CP-LC-0729 (50%) MC3] | (+) CP-LC-0729 /MC3/DOPE/Chol/DMG-PEG/DiR | 50/11.9/11.9/23.8/2.4/0 | 10 |
| [ (+) CP-LC-0729 (30%) CP-LC-0729] | (+) CP-LC-0729 / CP-LC-0729 /DOPE/Chol/DMG-PEG/DiR | 30/16.7/16.7/33.3/3.4/0 | 10 |
| [ (+) CP-LC-0729 (40%) CP-LC-0729] | (+) CP-LC-0729 / CP-LC-0729 /DOPE/Chol/DMG-PEG/DiR | 40/14.3/14.3/28.6/2.9/0 | 10 |
| [ (+) CP-LC-0729 (50%) CP-LC-0729] | (+) CP-LC-0729 / CP-LC-0729 /DOPE/Chol/DMG-PEG/DiR | 50/11.9/11.9/23.8/2.4/0 | 10 |
| [ (+) CP-LC-0729 (50%) CP-LC-0729]-DiR | (+) CP-LC-0729 / CP-LC-0729 /DOPE/Chol/DMG-PEG/DiR | 49.9/11.8/11.8/23.7/2.4/0.4 | 10 |

**Table S6.** Characterization of formulated LNPs by microfluidics for lung targeting strategy.

|  | Particle  size (nm) | P.D.I. | Zeta potential (mV) | Apparent  pKa | %EE |
| --- | --- | --- | --- | --- | --- |
| [DOTAP (50%) MC3] LNP | 58.68 | 0.1975 | 14.71 | 7.29 | 98.84 |
| [DOTAP (50%) CP-LC-0729] | 66.6 | 0.25 | 12.15 | 6.16 | 98.9 |
| [ (+) CP-LC-0729 (50%) MC3] | 78.6 | 0.37 | 7.13 | 8.16 | 99.2 |
| [ (+) CP-LC-0729 (30%) CP-LC-0729] | 65.9 | 0.26 | 6.48 | 7.83 | 98.6 |
| [ (+) CP-LC-0729 (40%) CP-LC-0729] | 81.9 | 0.35 | 9.366 | 7.93 | 99.1 |
| [ (+) CP-LC-0729 (50%) CP-LC-0729] | 71.7 | 0.30 | 11.54 | 7.37 | 99.2 |
| [ (+) CP-LC-0729 (50%) CP-LC-0729]d | 60.9 | 0.23 | 13.65 | 7.76 | 99.6 |

LNPs were formulated by microfluidic mixing. The resulting lipid particles encapsulating mRNAs were dialyzed overnight against a pH 8 Tris buffer solution containing 15% sucrose. Lipid particles were characterized by measuring their size distribution, polydispersity and Z-potential by dynamic light scattering (DLS). mRNA encapsulation was assessed by Quant-IT® Ribogreen following the manufacturer’s instructions. Apparent pKa was calculated using a 6-(p-toluidinyl) naphthalene-2-sulfonic acid (TNS) assay.

**Table S7.** Characterization and in vivo performance of formulated LNPs of lipid CP-LC-0729 in different lipid molar ratios by microfluidics.

| Lipid Molar Ratios  (CP-LC-0729/DOPE/Chol/DMG-PEG) | Size (nm) | PDI | Zeta potential (mV) | EE (%) | Total Flux (p/s) |
| --- | --- | --- | --- | --- | --- |
| 40/16/40/4 | 67.4 | 0.2679 | -3.135 | 98.0 | 7.15E+07 |
| 35.7/28.6/28.6/7.1 | 57.5 | 0.1544 | -7.321 | 91.0 | 2.21E+07 |
| 27.6/33.1/38.7/0.6 | 122.6 | 0.1760 | -9.176 | 96.8 | 8.34E+08 |
| 26/41.7/31.3/1 | 108.8 | 0.2087 | -6.754 | 96.3 | 2.45E+08 |
| 43.8/12.5/37.5/6.3 | 62.34 | 0.2737 | -6.922 | 93.4 | 3.11E+07 |
| 37.8/21.6/37.8/2.7 | 88.73 | 0.3088 | -12.28 | 95.8 | 1.45E+08 |
| 40.7/34.9/23.3/1.2 | 103.4 | 0.2180 | -8.246 | 95.7 | 8.50E+08 |
| 34.8/39.8/24.9/0.5 | 151.2 | 0.0571 | -6.022 | 94.7 | 1.03E+09 |
| 42.9/28.6/23.8/4.8 | 75.89 | 0.3078 | -13.32 | 81.8 | 5.27E+07 |
| 41.9/37.2/18.6/2.3 | 85.5 | 0.2122 | -8.296 | 89.4 | 1.63E+08 |
| 47.8/26.1/21.7/4.3 | 74.55 | 0.1584 | -12.5 | 70.4 | 1.16E+08 |
| 46.8/34/17/2.1 | 97.23 | 0.2288 | -8.555 | 86.7 | 1.95E+08 |
| 50/10/38.5/1.5 | 98.47 | 0.2114 | -9.349 | 91.3 | 7.05E+08 |

LNPs were formulated by microfluidic mixing at N/P ratios of 6 (mol/mol). The resulting lipid particles encapsulating mRNAs were dialyzed overnight against a pH 8 Tris buffer solution containing 15% sucrose. Lipid particles were characterized by measuring their size distribution, polydispersity and Z-potential by dynamic light scattering (DLS). mRNA encapsulation was assessed by Quant-IT® Ribogreen following the manufacturer’s instructions. Apparent pKa was calculated using a 6-(p-toluidinyl) naphthalene-2-sulfonic acid (TNS) assay. Mice were i.m injected with mRNA-Luc-loaded LNPs at an mRNA dose of 0.05 mg/kg. Luminescence imaging acquisition was performed at 4 h post-treatment and total flux was quantified. In vivo mLuc expression (n= 3 biologically independent samples). Data are presented as mean values.
